# A non-invasive, quantitative study of broadband spectral responses in human visual cortex

**DOI:** 10.1101/108993

**Authors:** Eline R. Kupers, Helena X. Wang, Kaoru Amano, Kendrick N. Kay, David J. Heeger, Jonathan Winawer

## Abstract

Currently, non-invasive methods for studying the human brain do not reliably measure signals that depend on the rate of action potentials (spikes) in a neural population, independent of other responses such as hemodynamic coupling (functional magnetic resonance imaging) and subthreshold neuronal synchrony (oscillations and event-related potentials). In contrast, invasive methods - animal microelectrode recordings and human intracortical recordings (electrocorticography, or ECoG) - have recently measured broadband power elevation spanning 50-200 Hz in electrical fields generated by neuronal activity as a proxy for the locally averaged spike rates. Here, we sought to detect and quantify stimulus-related broadband responses using a non-invasive method - magnetoencephalography (MEG) - in individual subjects. Because extracranial measurements like MEG have multiple global noise sources and a relatively low signal-to-noise ratio, we developed an automated denoising technique, adapted from Kay et al, 2013 (1), that helps reveal the broadband signal of interest. Subjects viewed 12-Hz contrast-reversing patterns in the left, right, or bilateral visual field. Sensor time series were separated into an evoked component (12-Hz amplitude) and a broadband component (60–150 Hz, excluding stimulus harmonics). In all subjects, denoised broadband responses were reliably measured in sensors over occipital cortex. The spatial pattern of the broadband measure depended on the stimulus, with greater broadband power in sensors contralateral to the stimulus. Because we obtain reliable broadband estimates with relatively short experiments (~20 minutes), with a sufficient signal-to-noise-ratio to distinguish responses to different stimuli, we conclude that MEG broadband signals, denoised with our method, offer a practical, non-invasive means for characterizing spike-rate-dependent neural activity for a wide range of scientific questions about human brain function.

**Author Summary:** Neuronal activity causes perturbations in nearby electrical fields. These perturbations can be measured non-invasively in the living human brain using electro- and magneto-encephalography (EEG and MEG). These two techniques have generally emphasized two kinds of measurements: oscillations and event-related responses, both of which reflect synchronous activity from large populations of neurons. A third type of signal, a stimulus-related increase in power spanning a wide range of frequencies (‘broadband’), is routinely measured in invasive recordings in animals and pre-surgical patients with implanted electrodes, but not with MEG and EEG. This broadband response is of great interest because unlike oscillations and event-related responses, it is correlated with neuronal spike rates. Here we report quantitative, spatially specific measurements of broadband fields in individual human subjects using MEG. These results demonstrate that a spike-rate-dependent measure of brain activity can be obtained non-invasively from the living human brain, and is suitable for investigating a wide range of questions about spiking activity in the human brain.

## 1 Introduction

The time-varying electric and magnetic fields near neural tissue provide an indirect but rich source of information about the activity of neural populations (<u>reviewed by 2</u>). These signals include rapid, ‘evoked’ responses that are time-locked to stimulus events (3), oscillatory responses (4), and nonoscillatory, broadband signals (5, 6). Broadband signals associated with sensory or motor tasks have been widely observed in human electrocorticography, or ‘ECoG’, (7) and animal microelectrode recordings (8). The broadband signal is an elevation in spectral power, typically spanning 50 to >200 Hz (9), and has attracted a great deal of attention for several reasons.

First, the broadband signal is correlated with the level of neural activity (multi-unit spiking), and hence provides a way to study population-level spiking activity in a cortical region (10-12). Second, the broadband signal has a smaller point spread function on the cortical surface than low frequency oscillations (8-25 Hz) (6, 13), and is therefore useful both for characterizing local properties of cortex and as a tool for neural prosthetics (14). Third, the broadband signal is correlated with a portion of the fMRI response and, together with other field potential measures, can be used to understand neural factors underlying an observed BOLD response (13, 15). Finally, because it can be measured at high temporal resolution, the broadband signal is useful for characterizing the temporal dynamics of neuronal activity (16, 17).

In contrast to intracranial recordings, in the extracranial measures of electroencephalography (EEG) and magnetoencephalography (MEG), broadband responses have not been widely and reliably observed. One significant challenge in identifying broadband in extracranial measures is that non-neural noise sources, particularly from miniature saccades, can be confounded with experimental designs, making neurally induced broadband responses hard to isolate (18-21).

A second challenge in measuring broadband extracranially is that the response is most evident in high frequencies (> 60 Hz), and the signal amplitude at these frequencies is low. While intracranial recordings have relatively high signal-to-noise ratios (SNR) even at these higher frequencies (7), EEG and MEG do not (22). Broadband signals can extend to lower frequencies (23, 24), but oscillatory processes in lower frequency bands often mask broadband measures in this range (6).

A third challenge is the potential confound between broadband signals and narrowband gamma oscillations. Narrowband gamma oscillations have been successfully measured with MEG and EEG, particularly in visual cortex for high contrast gratings (25-27). The frequency range of these oscillations (30-100 Hz) overlaps the broadband range, but the narrowband and broadband signals reflect biologically different processes (7-9, 11). The ability to measure one does not imply the ability to measure the other.

Here, we sought to measure broadband signals quantitatively in the human brain using a noninvasive method (MEG). In order for this important, spike-dependent signal to be useful, it is necessary to measure it reliably in individual subjects, with a high SNR. A high SNR is essential if this signal will be widely used to study differences across stimuli, tasks, or groups. We developed a novel, automated MEG denoising algorithm adapted from prior fMRI work (1). Our experiments were designed to elicit spatially localized neural responses in visual cortex, and eye movements were measured in a subset of subjects to test for possible confounds from non-neural sources.

## 2 Results

A large field ‘on-off’ stimulation experiment (Figure 1) was used to characterize the stimulus-locked (steady state evoked field, ‘SSVEF’) and broadband responses in visual cortex measured with MEG. The two measures are reported below, both prior to and after applying our new denoising algorithm.

**Figure 1.**
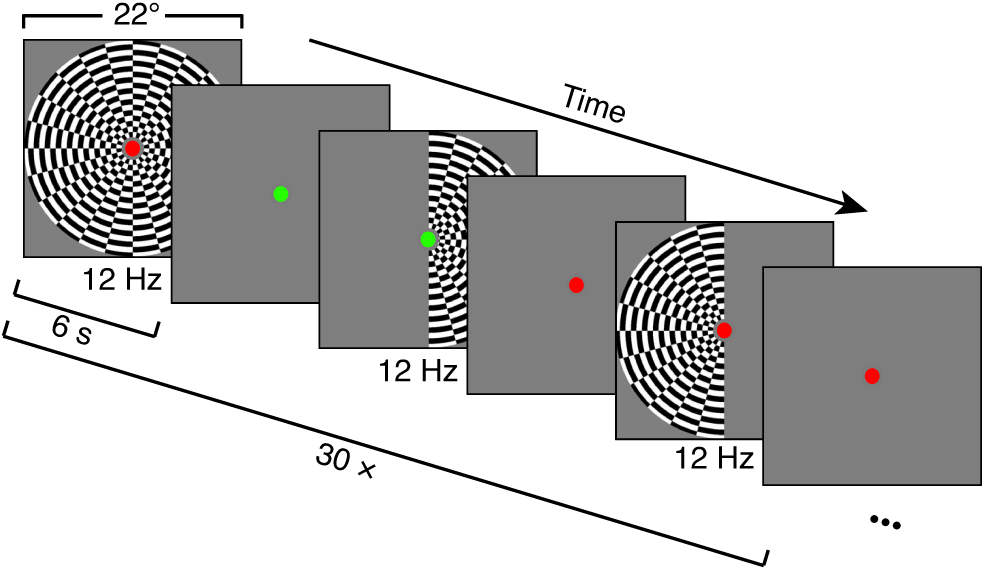
Overview of experimental design. Large-field on-off stimuli were presented in 6-s blocks consisting of either both-, left-, or right-hemifield contrast reversals (12 reversals per second), alternating with 6-s blocks of blanks (mean luminance). A run consisted of six stimulus blocks and six baseline blocks, after which the subject had a short break. The figure shows the first half of one run. Within a run, the order of both-, left-, and right-field contrast reversing periods was randomized. Fifteen runs were obtained per subject, so that there were 30 repetitions of each stimulus type across the 15 runs. The fixation dot is increased in size for visibility. Actual fixation dot was 0.17 degrees in radius (6 pixels).

### 2.1 Stimulus-locked and broadband signals measured with MEG

In each stimulus condition (left-, right-, and both-hemifield), the stimulus contrast reversed 12 times per second, so a stimulus-locked signal was measured at 12 Hz and harmonics. For the purposes of summarizing responses, we binned the data into non-overlapping 1-second epochs, and discarded the first of 6 epochs per block (see Methods). Because the largest component was at 12 Hz, we defined the stimulus-locked signal for a particular stimulus condition as the amplitude at 12 Hz, averaged over all 1-second epochs with that stimulus (typically ~180 epochs) computed for each of the 157 sensors in each subject (Figure 2A). The broadband signal was computed by averaging the log power across frequencies between 60 and 150 Hz, excluding multiples of the stimulus frequency (12 Hz), and then exponentiating the mean (Figure 2B). For both stimulus-locked and broadband responses, we defined the *signal* for a given stimulus condition as the median of the bootstrapped difference between the response to stimulus and blank, and the variability, or *noise,* as half the 68% confidence interval of this distribution (Figure 2B, inset); Hence the *SNR* was the quotient of these two numbers.

**Figure 2.**
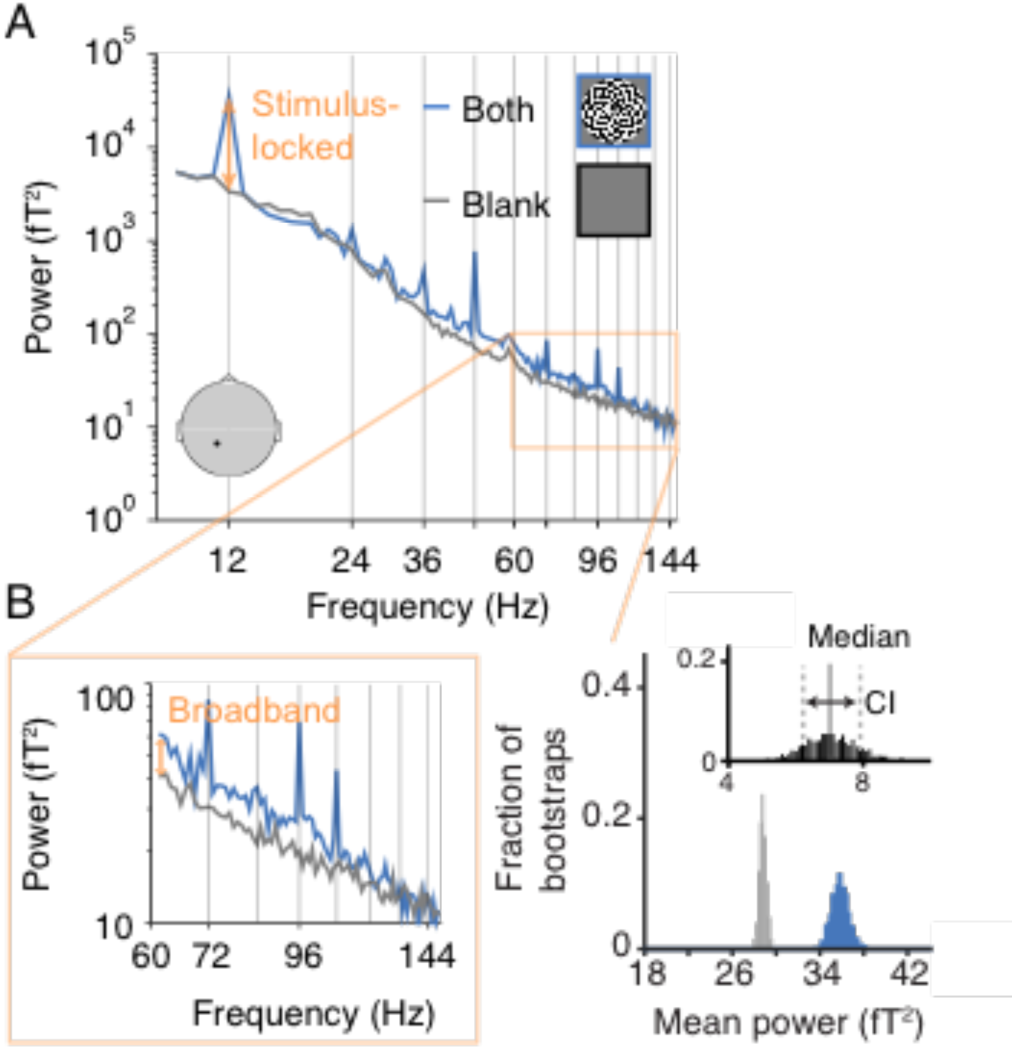
Example response to flickering large-field stimulus. **(A)** Spectral power, averaged across 180 1-second epochs, during which the subject viewed either the both-hemifield stimulus (blue line) or a blank screen at mean luminance (gray line). The black dot on the schematic head indicates the location of the sensor. The peak at 12 Hz corresponds to the frequency of dartboard contrast reversals, and is a measure of the stimulus-locked component (orange arrow). **(B)** The left plot zooms in on higher frequencies to emphasize the broadband component, most evident in this example data set as a spectral power elevation spanning 60 to 150 Hz. The increase in the broadband response of the stimulus condition relative to the blank condition is shown by the orange arrow. The histograms on the right show the broadband level separately for the stimulus condition (blue) and the blank condition (gray), and the difference between them (black), computed 1000 times by bootstrapping over epochs in the experiment. The SNR was defined as the signal, the median of the difference distribution, divided by the noise, half the 68% confidence interval. Data from subject S1. Made with function *dfdMakeFigure2.m.*

Both the stimulus-locked and broadband signals were largest in medial, posterior sensors, as expected from activations in visual cortex (28). For the stimulus-locked signal, the both-hemifield condition tended to produce broadband signals in bilateral posterior sensors, whereas the single-hemifield conditions produced responses that were lateralized, with higher SNR contralateral to the stimulus. This pattern could be seen in an example subject and in the average across subjects (Figure 3). The lateralization of the stimulus-locked signal was less clear in the average across subjects due to imperfect alignment of the sensors showing the largest differential response to the left- and right-hemifield stimuli. In each of the 8 individual subjects and in each of the 3 conditions, the stimulus-locked response was evident, with the signal at least 10x above the noise (Supplementary Figure 1A).

**Figure 3.**
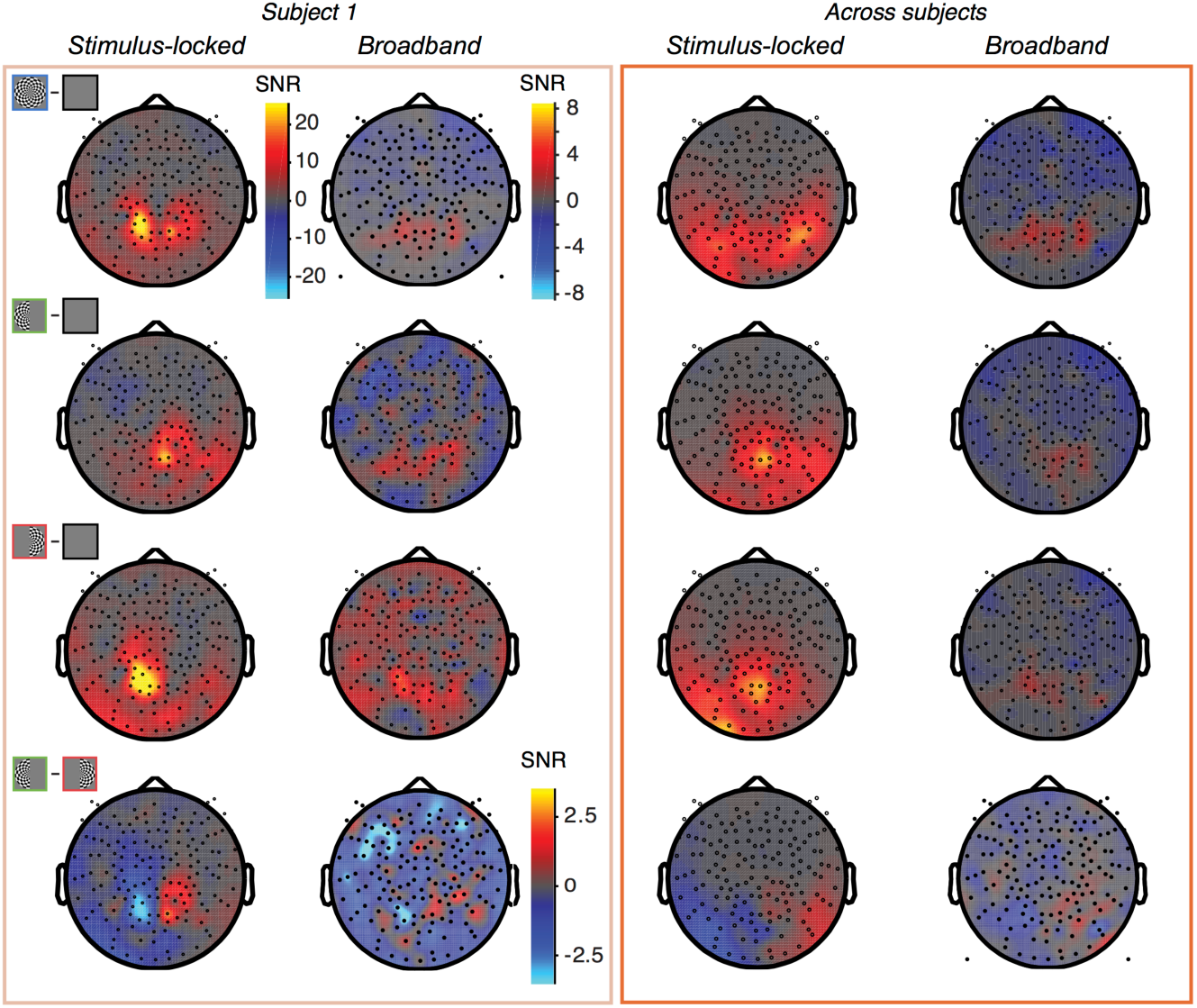
Topographic map of stimulus-locked and broadband responses. Data from subject S1 **(left)** and averaged across subjects S1-S8 by sensor **(right).** The top 3 rows show data from the 3 stimulus conditions (both-, left-, and right-hemifield) compared to blank, and the lower row shows data as the left-only minus right-only conditions. The dependent variable plotted for the single subject data is the signal-to-noise ratio at each sensor, computed as the mean of the contrast (stimulus minus blank) across bootstraps divided by the standard deviation across bootstraps (bootstrapped over epochs). For the group data, the signal-to-noise ratio is the mean of the subject-specific SNRs at each given sensor. The same scale bar is used for all stimulus-locked plots. For the broadband plots, one scale bar is used for the first three rows, and a different scale bar with a smaller range is used for the fourth row. Made with *dfdMakeFigure3.m.*

The spatial pattern of broadband signals was qualitatively similar to the spatial pattern of the stimulus-locked signal, with bilateral posterior responses in the both-hemifield condition, and lateralized responses in the single-hemifield conditions (Figure 3, individual example and group-averaged data). However, the broadband responses had much lower signal-to-noise than the stimulus-locked responses, and in many of the individual subjects, broadband was not evident in one or more conditions; for example, in the both-hemifield condition, there was a clear medial posterior broadband response for S1, S2, and S3, with signal at least 5x above noise, but not for S4 (Supplementary Figure 1B). The broadband responses were less reliable for the left- and right-hemifield conditions than for the both-hemifield conditions.

The fact that broadband responses were evident in a few subjects in some conditions indicates that it is possible to measure broadband fields with MEG. However, if this signal cannot be measured reliably in many subjects and many conditions, then the practical value of measuring broadband with MEG is limited. This motivated us to ask whether denoising the MEG data could unmask broadband signals, making it more reliable across subjects and stimulus conditions.

### 2.2 Denoising increases the broadband SNR by reducing variability

The MEG data were denoised using a new algorithm as described in detail in the Methods section. In brief, for each subject a subset of sensors that contained little to no stimulus-locked responses were defined as the noise pool (Figure 4A). Once the noise-pool was defined, the time series in each sensor and in each epoch was filtered to remove all signals not contributing to the broadband measurement (Figure 4B). Global noise regressors were then derived by principal component analysis from the filtered time series in the noise pool in each 1-s epoch (Figure 4C). The first 10 PCs were projected out of the data in each sensor, epoch by epoch (Figure 4D). Subsequently, the broadband responses were defined the same way as in the non-denoised data set (Figure 2B).

**Figure 4.**
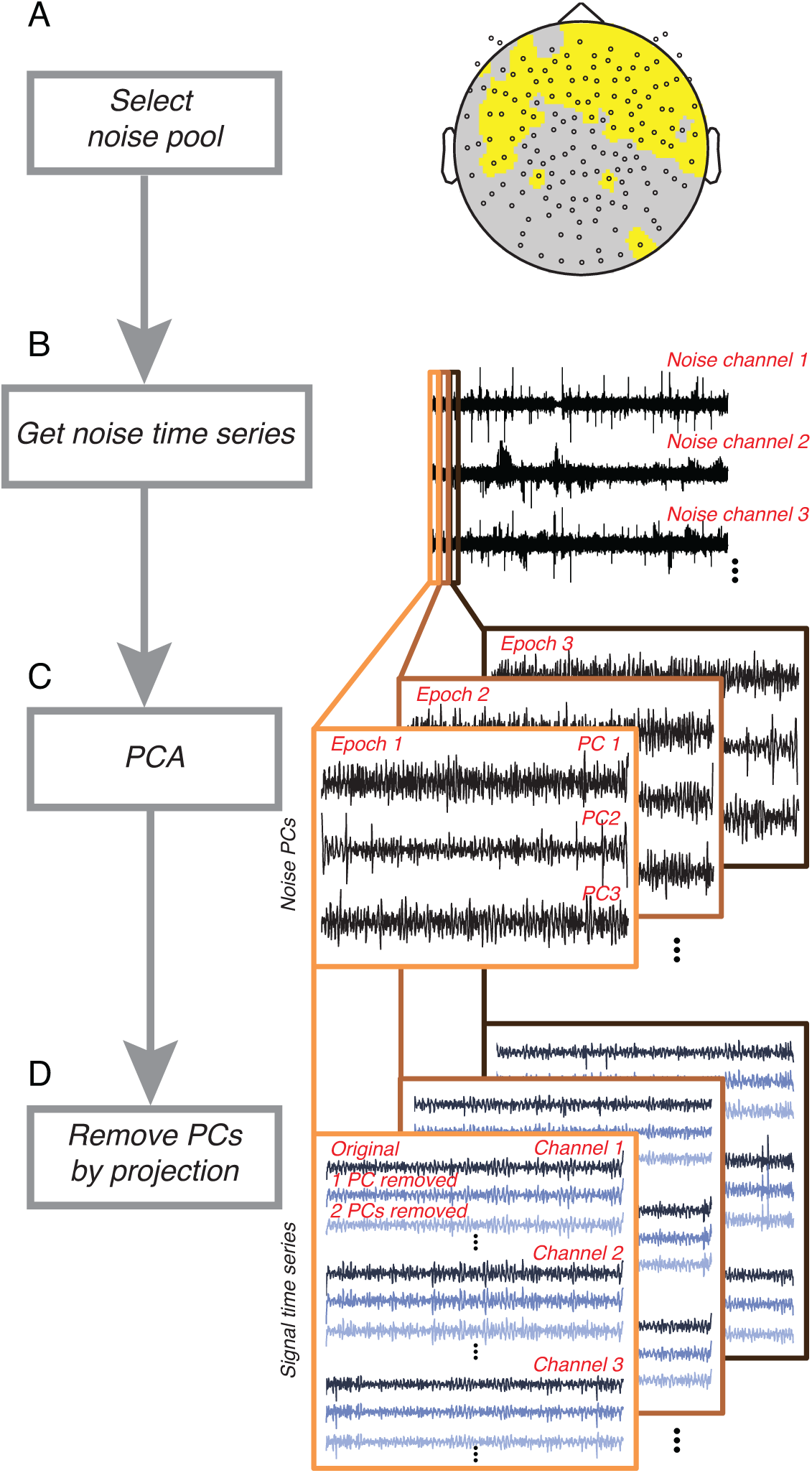
Denoising procedure. Following an estimate of response reliability computed from non-denoised data (Figure 2), the algorithm first selects a noise pool. **(A)** The noise pool is comprised of sensors whose SNR from the evoked (stimulus-locked) component falls below a threshold. **(B)** The time series from each sensor in the noise pool is then filtered to remove components that do not contribute to the broadband computation. **(C)** Principal component analysis is then computed within each epoch. **(D)** for each epoch, the first *n* PCs are projected out from the time series of all sensors, yielding *n* new data sets. For each new data set, broadband responses were recomputed, as in Figure 2B.

We first illustrate the effect of denoising with an example from a single sensor in one subject (Figure 5). This sensor showed a broadband response both prior to, and after, denoising. The benefit of denoising was not evident when comparing the mean power spectra before and after denoising (Figure 5A). Denoising did not reduce the variability in power across frequencies, nor did it increase the separation in the spectra for the contrast stimulus and the blank. Instead, the effects of denoising are better appreciated by examining the variability across epochs rather than across frequencies (Figure 5B). The biggest effect is that the broadband power estimates became less variable across epochs, both for the blank condition and the stimulus condition. This is indicated by the narrower distributions in the response amplitudes for the two conditions (Figure 5B, main panels) and for the difference between conditions (Figure 5B, insets). The standard deviation of the difference distributions decreased more than two-fold (from 0.79 to 0.35) as a result of denoising.

**Figure 5.**
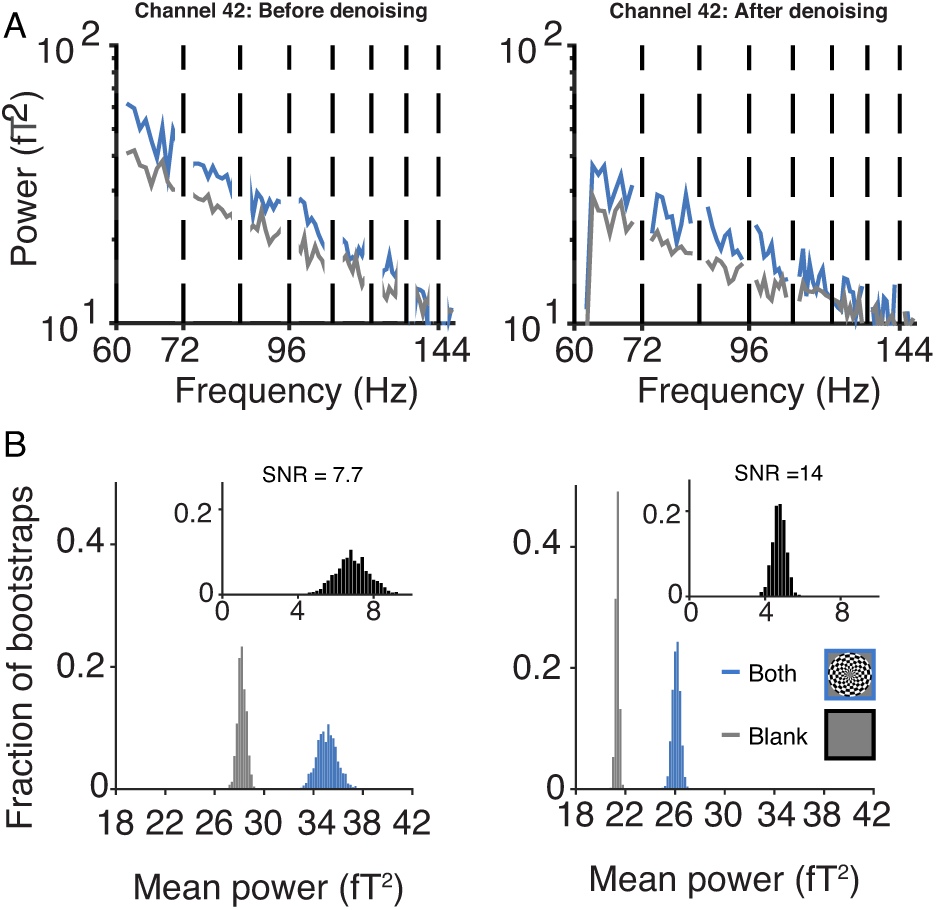
Effect of denoising on broadband response. **(A)** The upper panel shows the power spectra from sensor 42, subject 1, averaged across 178 epochs with the both-hemifield stimulus (blue) and blank screen (gray). The left panel is prior to denoising and is identical to the inset in Figure 4, except that harmonics of stimulus-locked frequencies have been removed. The right panel is the same as the left, except after denoising. **(B)** The lower panel shows the distributions of the bootstrapped broadband power for the both-hemifield (blue), blank (gray), and both-hemifield minus blank (black, inset), prior to denoising (left) and after denoising (right). The SNR is defined as the median of the difference distribution divided by half of the 68% confidence interval in the difference distribution (7.7 prior to denoising, 14.0 after). The effects of denoising are to reduce the mean power, and more importantly, reduce the standard deviation across epochs. Made with *dfdMakeFigure5.m.*

There are two other secondary patterns evident in these distributions. First, the mean broadband power of both the blank and stimulus condition decreased as a result of denoising (for the both-hemifield condition, 35.8 versus 26.1, prior to versus after denoising; for the blank, 28.7 versus 21.4). This was expected because projecting out signal reduces power. Second, the contrast between the two conditions (difference between the means) reduced: 7.0 prior to denoising versus 4.8 after denoising. The combination of these two effects was that the *percent difference* was little changed, with broadband power from the contrast-stimulus about 25% more than for the blank before and after denoising. Hence denoising did not increase the estimate of the percent signal change.

It is important to consider how these effects interact. Because the reduction in variability across epochs was the biggest effect of denoising (more than 2-fold), there was more than a doubling of SNR, computed as the median divided by the variability of the difference distribution. In sum, the spectral plots show that the variability in power *across frequencies* was little affected by denoising (Figure 5A), whereas the distribution plots show that the variability in total broadband power *across epochs* was reduced considerably (Figure 5B).

We now consider the effect of denoising across sensors, subjects, and stimulus conditions. Projecting out noise PCs substantially increased the signal-to-noise ratio of the broadband measurement in visually responsive sensors. For example, in the both-hemifield condition for subject S1, the median SNR of the 10 most visually responsive sensors increased from 5 to 10 after denoising (Figure 6A, blue solid line), similar to the example sensor shown earlier (Figure 5B). In contrast, the SNR of the 75 sensors in the noise pool was relatively unaffected by denoising (Figure 6A, blue dashed line). This was expected because sensors in the noise pool were unlikely to distinguish stimulus from blank. Across the 8 subjects in the both-hemifield condition, taking the mean of the 10 most visually responsive sensors for each subject, the SNR increased about 3-fold (from 1.6 to 5.0), with a numerical increase in every subject (Figure 6B). Because the SNR stabilized 11 in all subjects with 10 or fewer PCs projected out, in subsequent analyses, for simplicity we report the effects of denoising with exactly 10 PCs. A comparison of the SNR before denoising (0 PCs projected out) and after (10 PCs projected out) summarized across all subjects and the three stimulus conditions shows increases in SNR for every subject in all conditions (Figure 6C) (*p*=0.0001, *p*=0.0007, *p*=0.0022 for two-tailed t-tests, 0 v 10 PCs, for both-, left-, and right-hemifield conditions, respectively).

**Figure 6.**
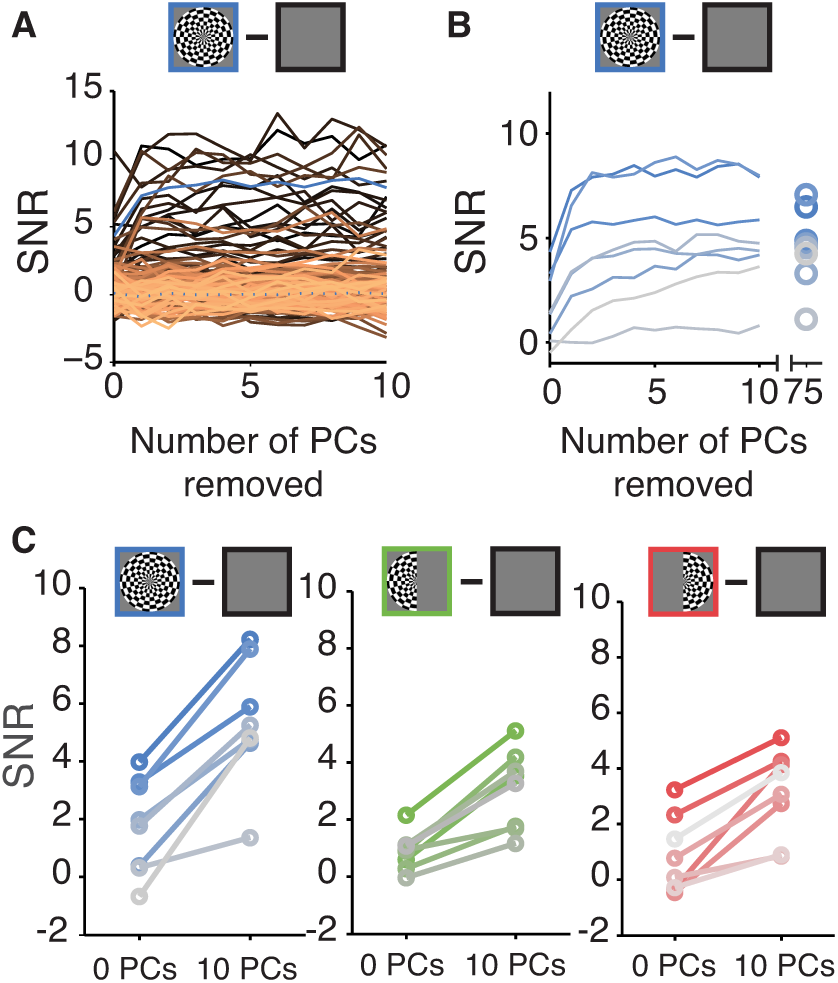
Effect of denoising on broadband SNR. **(A)** SNR as a function of the number of PCs projected out in subject S1 for the both-hemifield stimulus. Each line is one sensor. The heavy blue line is the mean of the 10 sensors with the highest SNR, as measured either before or after denoising. **(B)** SNR as a function of PCs projected out in each of 8 subjects for the both-hemifield stimulus. Each line is the mean across the 10 sensors with the highest SNR in one subject. The rightmost points indicate the effect of projecting out all 75 PCs. **(C)** SNR before denoising (0 PCs projected out) and after denoising (10 PCs projected out) for each stimulus condition. Each line is the mean of the 10 sensors with the highest SNR for one subject in one stimulus condition. Color saturation corresponds to the subject number (highest to lowest saturation, subjects 1-8, respectively). Made with *dfdMakeFigure6.m.*

In principle, the SNR increases could have arisen from increased signal, decreased noise, or both. To distinguish among these possibilities, we compared the signal level alone and the noise level alone before and after denoising. As in prior results, the signal was defined as the difference in broadband power between the contrast pattern and the blank (median across bootstraps), and the noise was defined as the variability of this difference metric (half of the 68% confidence interval across bootstraps). For all three stimulus conditions in most subjects, the signal was largely unaffected by denoising, staying at a similar level or decreasing slightly, while the noise level went down substantially (Figure 7). These analyses indicate that the increase in SNR from denoising (Figure 6) was caused by a reduction in epoch-to-epoch variability of the broadband signal level, and not by an increase in the signal level, consistent with the results of the single example sensor (Figure 5). Expressed as a percentage increase over baseline, the broadband response to the both-hemifield stimulus after denoising was ~10.9±1.7% averaged across the top 10 sensors in each subject (mean ± sem across subject), and 12.6%±1.6% for the top 5 sensors. This contrasts with the much larger stimulus-locked response, which was a nearly 8-fold increase over baseline even prior to denoising (678%±226% increase over baseline for the top 5 sensors).

**Figure 7.**
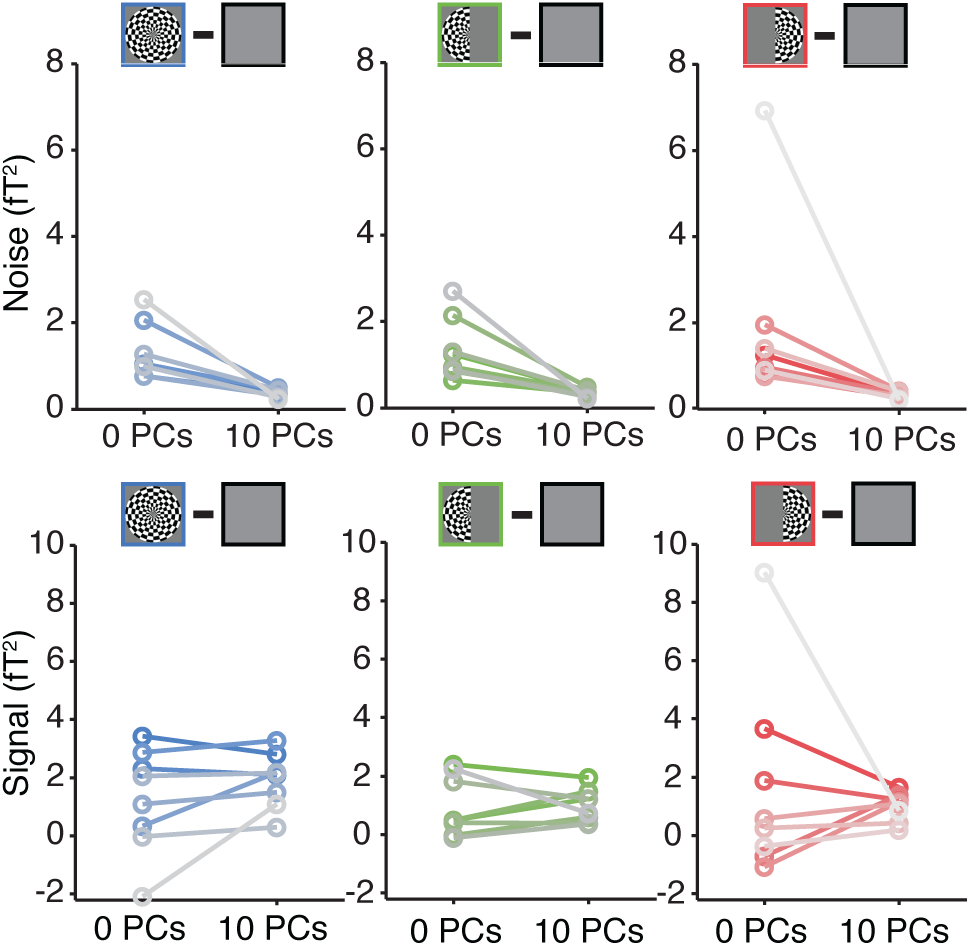
Effect of denoising on the broadband signal and noise. Noise **(upper)** and signal **(lower)** before and after denoising in each of three stimulus conditions. Plotting conventions as in Figure 6c. Made with *dfdMakeFigure7.m.*

The effect of denoising the broadband signal was not uniform across the sensor array. In general, sensors where we expected visual activity (over the posterior, central part of the head) showed increased SNR following denoising. In the example subject S1 as well as the average across subjects, the denoised broadband response was observed in bilateral sensors for the both-hemifield condition, and with a contralateral bias (relative to the midline) in the two lateralized conditions (Figure 8). For the both-hemifield stimulus, broadband responses were evident in sensors over the posterior, middle of the head in most individual subjects (Figure 9).

**Figure 8.**
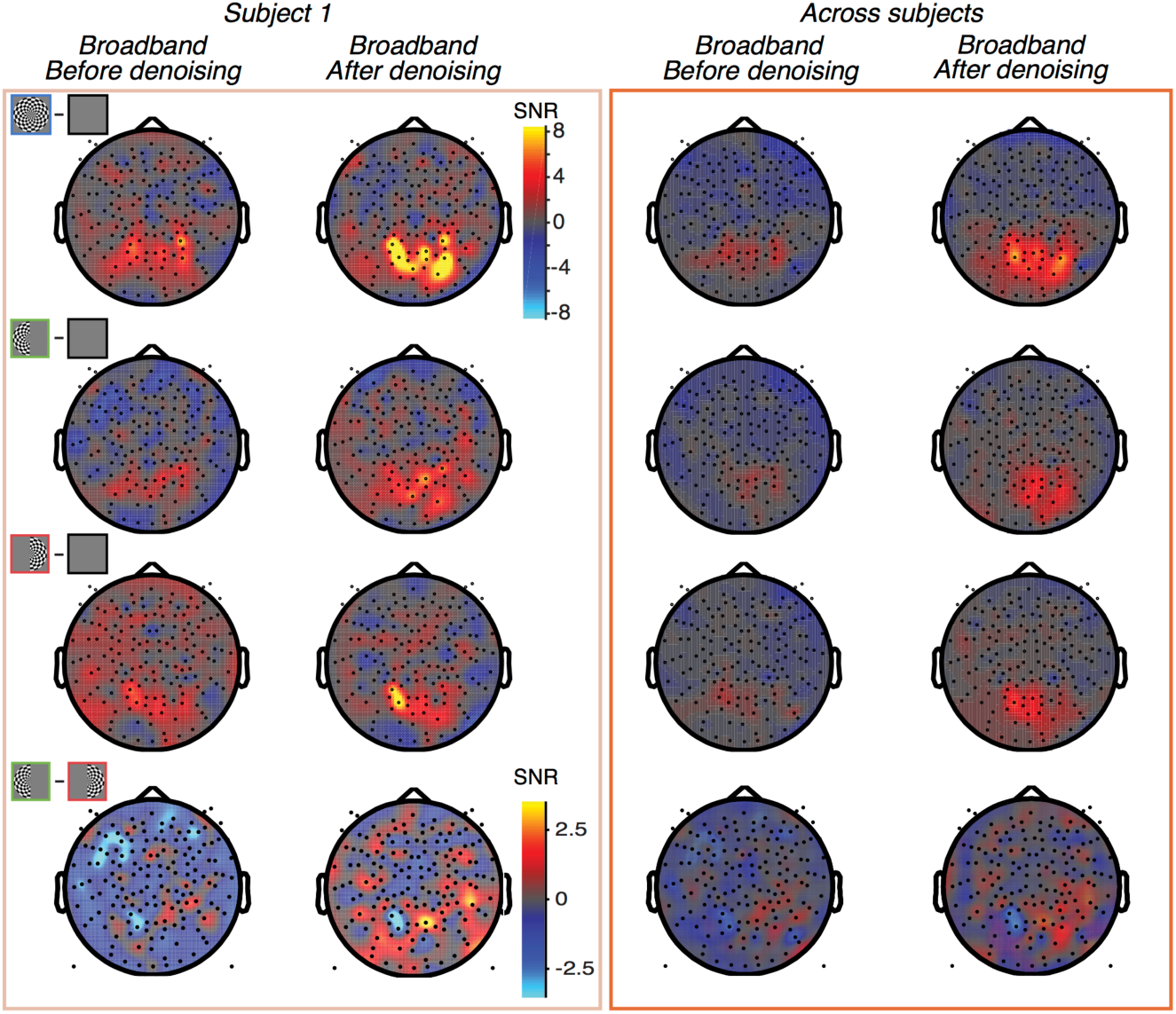
Topographic map of broadband SNR before and after denoising. Data from subject S1 **(left)** and averaged across subjects S1-S8 by sensor **(right).** The top 3 rows show data from the 3 stimulus conditions (both-, left-, and right-hemifield) and the fourth row shows the difference between the left-only and right-only conditions. The fourth row uses a different scale bar from the other 3 rows. The columns show data before and after denoising. Made with *dfdMakeFigure8. m.*

**Figure 9.**
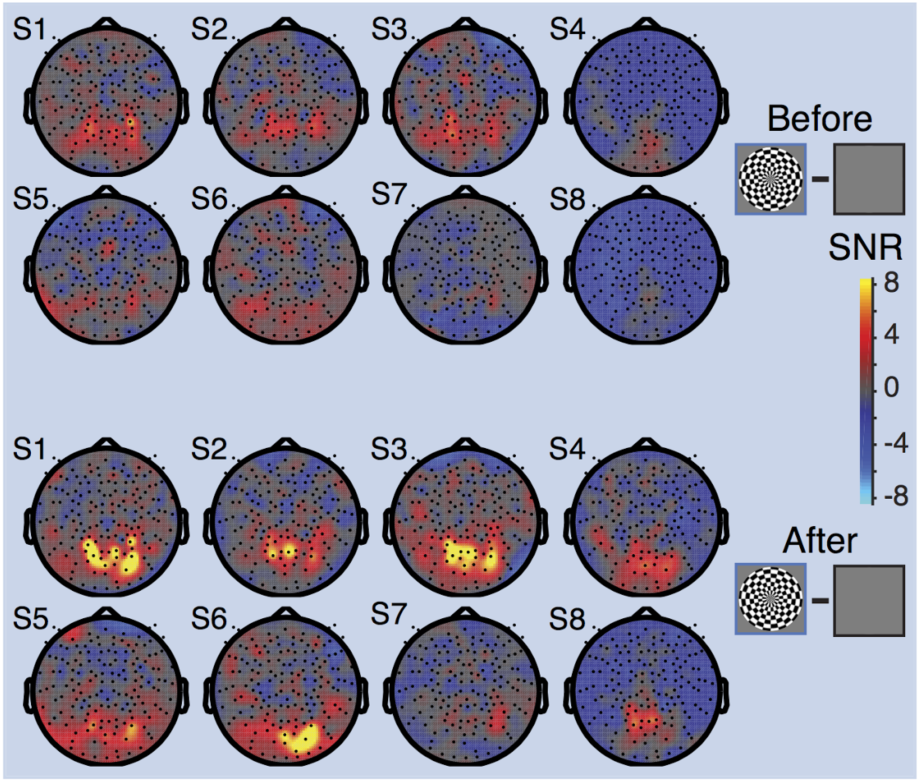
Topographic maps of broadband SNR in individual subjects after denoising. Head plots show the SNR for the both-hemifield stimulus, before denoising (**above**) and after denoising (**below**). For individual subject data for all stimulus conditions, see Supplementary Figure 2. Made with *dfdMakeFigure9.m.*

### 2.3 Control analyses for MEG Denoise algorithm

To validate the assumptions in our denoising algorithm, we ran three control analyses. In one control analysis, we concatenated all epochs to derive noise regressors from the whole experimental time series (Figure 10, 2^nd^ bar, compared to using the default of 1-s epochs to derive noise regressors - 1^st^ bar). The elevation in broadband SNR was significantly less when we concatenated all epochs (*p* = 0.0016, *p* = 0.0023 and *p* = 0.0447, for the three stimulus conditions respectively). In the second control analysis, the noise pool included all sensors rather than only those sensors that were not visually responsive. Here, the noise regressors included some signal as well as noise, and hence should be of less benefit. This expectation was confirmed, in that there was no increase in SNR when the algorithm was run with the omission of the noise-pool-selection step (Figure 10, 3^rd^ bar, *p* = 0.0014, *p* = 0.0015 and *p* = 0.0020 for the three stimulus conditions respectively). In a 3^rd^ control analysis, we phase-scrambled each of the epoch-by-epoch noise time series. The phase-scrambled regressors were temporally uncorrelated with the actual time series in the noise. As a result, we found no change in SNR levels (Figure 10, fourth bar, *p* = 0.0001, *p* = 0.0003 and *p* = 0.0017).

**Figure 10.**
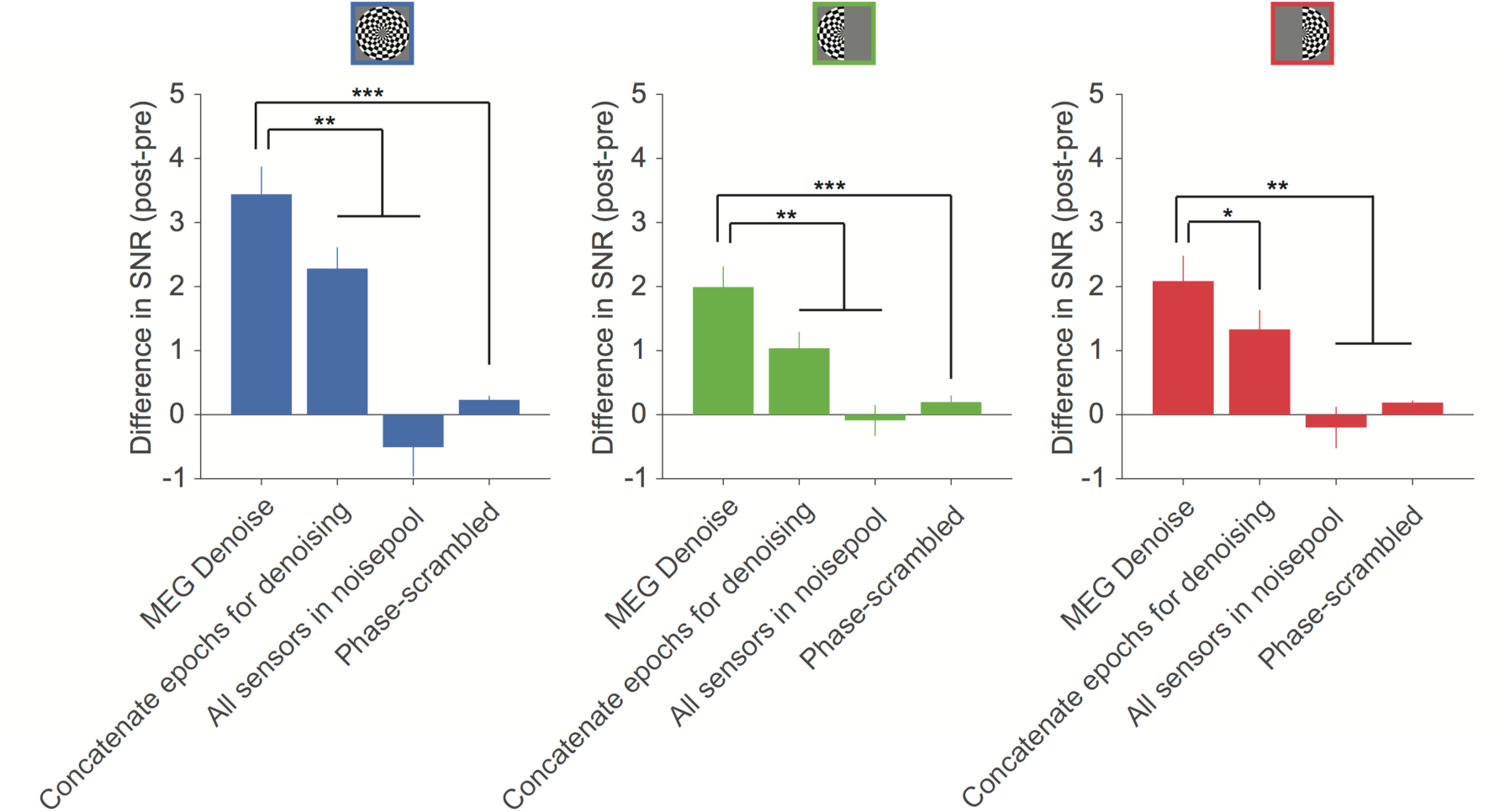
Comparison of MEG Denoise to control analyses. When the denoising algorithm derives noise regressors from the whole experimental time series (‘Concatenate epochs for denoising’), the amount of SNR gain is significantly less than the standard MEG Denoise (regressors derived separately from each 1-s epoch). When the noise regressors are derived from all sensors (‘All sensors in noisepool’), or when the time series of the regressors are phase-scrambled, there is little or no change in SNR for all three stimulus conditions. Statistical significance is computed by a 2-tailed t-test, paired by subject, between denoising analyses. Statistical significance is indicated by * = p < 0.05, ** = p < 0.01, *** = p < 0.001 between the MEG Denoise algorithm and each of the other controls. Made with *dfdMakeFigure10.m.*

### 2.4 Other denoising algorithms

To assess how other existing denoising algorithms affect our measurement of broadband power, and how they interact with our new denoising algorithm, we ran two different denoising algorithms, either alone or in combination with MEG Denoise. The two algorithms we tested were CALM, or continuously adjusted least-square method (29) and TSPCA, or time-shift principal component analysis (30). Both of these make use of reference MEG sensors which face away from the head and measure environmental rather than physiological fields. By design, these algorithms project out time series from the subspace spanned by the reference sensors, thereby reducing environmental noise, but not physiological noise. Applying either one of these two algorithms alone to the 8 data sets reported above increased the broadband signal-to-noise ratio, evident in the group-averaged sensor plots (Figure 11A, columns 3-4 versus column 2), and the increased SNR in the 10 most responsive sensors (Figure 11B, 2^nd^ and 3^rd^ bar versus 1^st^ bar in each plot).

**Figure 11.**
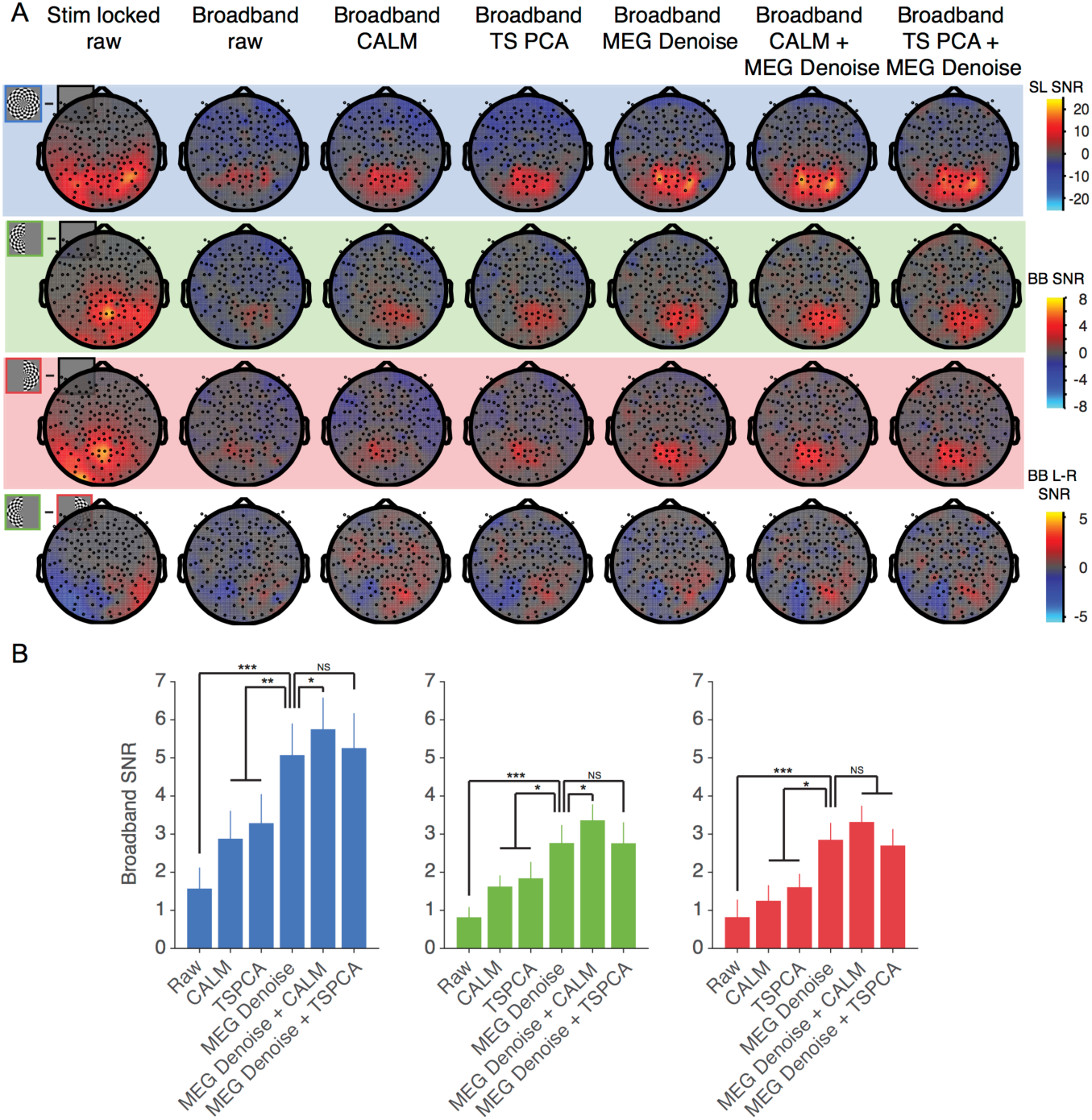
Comparison of different denoising algorithms on NYU datasets (averaged across subjects S1-S8). (A)The columns represent SNR values for the stimulus-locked signal (column 1), broadband signal without denoising (column 2), and broadband signal with one or more denoising algorithms. One scale bar is used for all stimulus-locked plots (column 1). A second scale bar is used for all broadband plots (columns 2-7) except for the Left minus Right plots (row 4, columns 2-7). Other details as in Figure 5. **(B)** Broadband SNR using different algorithms for both-hemifield (left), left-hemifield (center) and right-hemifield (right) stimuli. Each bar is the change in SNR from baseline (column 2 in panel A), averaged across the top 10 sensors per subject (mean +/ − SEM across subjects). Top sensors were defined as the 10 sensors from each subject with the highest SNR across any of the 3 stimulus conditions and any of the denoising algorithms (columns 2-7). Statistical significance computed and indicated as in Figure 10. Made with *dfdMakeFigure11.m.*

In planned comparisons, we evaluated the SNR increase of each algorithm or combination of algorithms to the increase from MEG Denoise alone. The increase from each of the two environmental algorithms alone was significantly less than that from our new MEG Denoise algorithm (Figure 11A, column 5 versus columns 3-4; Figure 11B, 4^th^ bar versus 2^nd^ and 3^rd^). Applying two algorithms in sequence, first either CALM or TSPCA, followed by MEG Denoise, also resulted in a large increase in broadband SNR (Figure 11A, columns 6 and 7). For all three stimulus conditions, the combination of MEG Denoise and CALM resulted in the largest gain in SNR, significantly larger than MEG Denoise alone for two out of the three conditions (Figure 11B, 5^th^ versus 4^th^ bars). This indicates that the MEG Denoise algorithm and an environmental algorithm captured some independent noise.

### 2.5 Effect of denoising on stimulus-locked SNR

In a separate analysis, we ran the MEG Denoise algorithm to evaluate its effect on the stimulus-locked signal. The methods were identical to those used to denoise the broadband signal except for the omission of one step, the step in which we filtered the time series to remove temporal components that do not contribute to the broadband signal. Denoising modestly increased the stimulus-locked SNR for all stimulus conditions for most subjects (Figure 12, top). The SNR increased numerically in all subjects (n=8) and in all stimulus conditions, although the percentage increases were smaller than those for denoising the broadband signal, ~20% increase compared to two-fold. As in the case of denoising the broadband signals, the main contribution to the increase in SNR for the stimulus-locked signal was a decrease in variability across epochs (Figure 12, bottom), rather than an increase in the signal level (Figure 12, middle).

**Figure 12.**
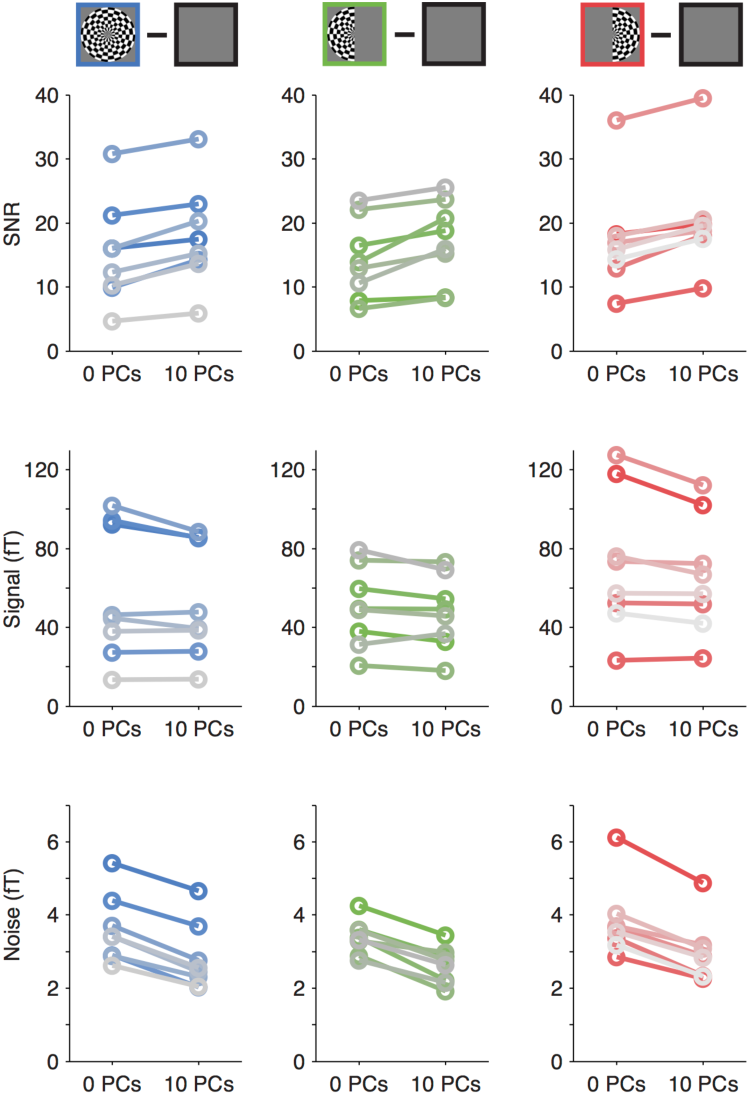
Denoising the stimulus-locked signal. The MEG Denoise algorithm results in a modest increase in SNR for most subjects in all three stimulus conditions **(top row).** This benefit is largely due to the fact that the noise level goes down from denoising **(middle bottom)** rather than the signal increasing **(middle row).** Plotting conventions as in Figure 7c and Figure 8. Made with *dfdMakeFigure12.m.*

### 2.6 Broadband fields measured with Elekta 360 Neuromag

To test whether the findings reported above generalize to other instruments and experimental environments, we conducted the same experiment using a different type of MEG system, an Elekta 360 Neuromag at CiNet. The CiNet system contains paired planar gradiometers, in contrast to the axial gradiometers used in the Yokogawa MEG at NYU, and the scanner is situated in a different physical environment, with potentially very different sources of environmental noise. The preprocessing pipeline at this imaging center often includes a denoising step based on temporally extended signal source separation (tSSS) (31, <u>32</u>). This additional experiment gave us the opportunity to ask several questions: (1) Are broadband fields observed with a different MEG sensor type and different physical environment? (2) Does the tSSS algorithm increase the broadband SNR? (3) Does our new MEG Denoise algorithm increase the SNR of data that have already been denoised with the tSSS algorithm?

The identical experiments were conducted with 4 new subjects. As expected, all three stimulus types led to a large stimulus-locked response in the posterior sensors, with a peak SNR of more than 10 in the group averaged data (Figure 13A, column 1). A modest, spatially specific broadband signal was measured from the undenoised data for each stimulus type (Figure 13A column 2), with a peak SNR of 1-2 in the group-average data for all three conditions. Unlike the NYU data, in the CiNet data the MEG Denoise algorithm on the raw data did not generally result in an increase in the broadband SNR (group data, Figure 13A, columns 2 and 3; individual subjects, Figure 13B, left side of each subplot). However, when the raw data were pre-processed with the tSSS algorithm (Figure 13A, column 4), application of MEG Denoise increased the SNR in all 3 stimulus conditions for 3 out of 4 subjects, and in 2 out of 3 stimulus conditions for the 4th subject. Together, the MEG Denoise algorithm increased the SNR by 2-3 fold, similar to the NYU data (both-hemifield: 2.8 to 5.6; left-hemifield: 0.8 to 2.4; right-hemifield: 2.01 to 4.4; means across subjects 1-4, top 10 sensors each, for the tSSS data and the MEG Denoised tSSS data). Just as with the NYU MEG data set, the combination of an algorithm tailored to find environmental noise (tSSS) and our algorithm produced the most robust results, indicating that MEG Denoise and the environmental denoising algorithm removed at least some independent sources of noise.

**Figure 13.**
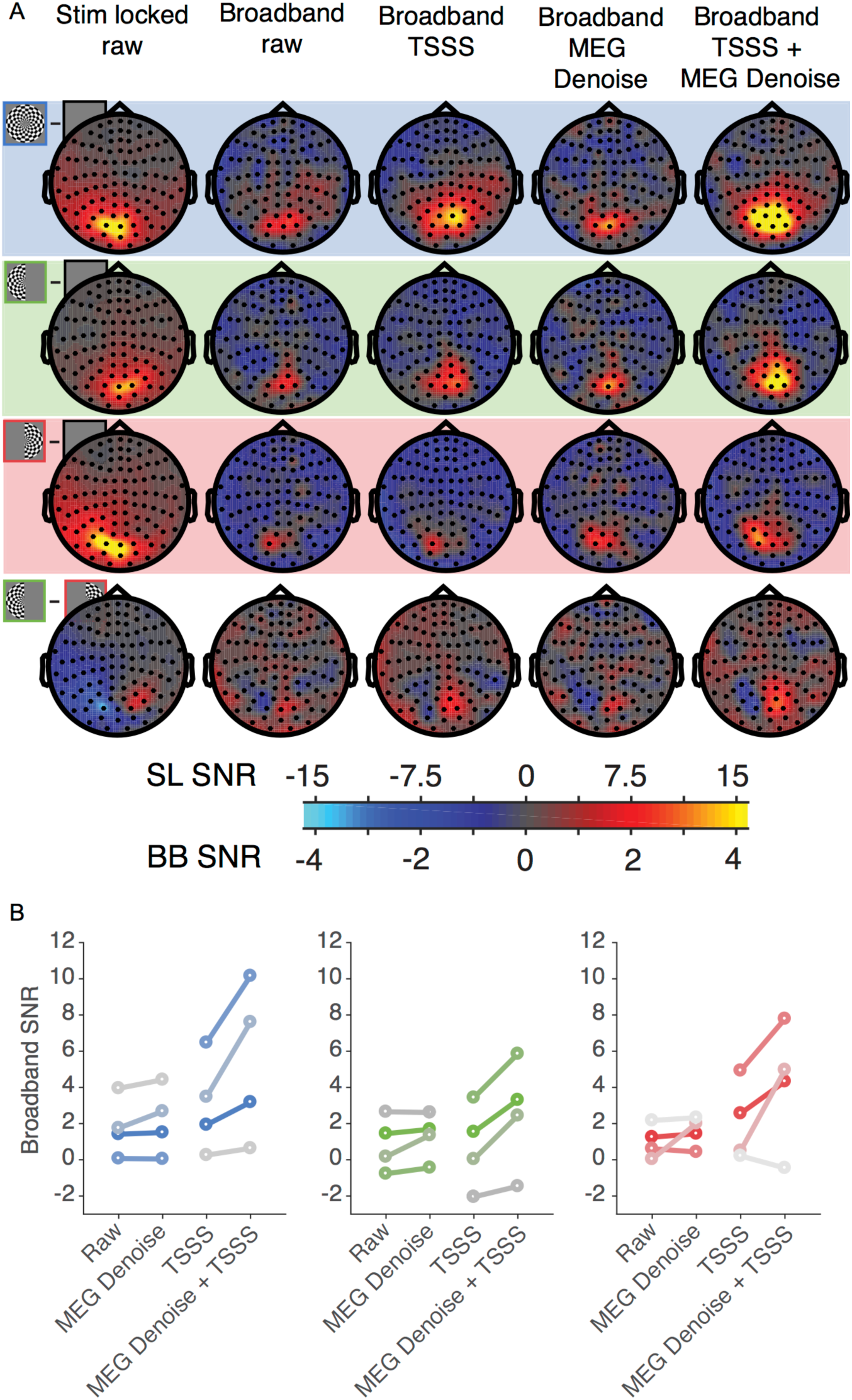
MEG data from CiNet Neuromag. **(A)** All plots show data averaged across new 4 subjects (S9-S13) in sensor space (sensor-wise mean of the subject SNR). The columns represent SNR values for the stimulus-locked signal (column 1), broadband signal without denoising (column 2), and broadband signal with one or more denoising algorithms. The same scale bar is used for all broadband data (columns 3 - 5). Other details as in Figure 3. **(B)** Broadband SNR using different algorithms for both-hemifield (left), left-hemifield (center) and right-hemifield (right) stimuli. Each line is average broadband SNR across the top 10 sensors for one individual. Top sensors were defined as the 10 sensors from each subject with the highest SNR across any of the 3 stimulus conditions and any of the denoising algorithms (columns 2-6). Made with *dfdMakeFigure13.m.*

### 2.7 Saccadic eye movements during MEG experiments

Saccadic eye movements are known to have a large influence on MEG and EEG measurements. This influence can be especially pernicious when measuring high frequency broadband signals, because the spike field (MEG) or spike potential (EEG) arising from extraocular muscle contraction can be spectrally broadband and can co-vary with task design; hence, it can easily be confused with broadband signals arising from brain activity (19, 21). For visual experiments, the spike potential in EEG is especially problematic because it tends to affect sensors which are also visually sensitive (posterior middle). In contrast, the MEG spike field is lateral, potentially influencing temporal and frontal sensors, with little to no effect on posterior sensors (18). Hence spike field artifacts are unlikely to contaminate our visually elicited broadband signals, which are most clearly evident in the central posterior sensors.

Nonetheless, for a subset of subjects (S6-S8), we measured eye movements during the MEG experiments and quantified the frequency of microsaccades, and the distribution of microsaccade direction, for each stimulus condition. Each of these 3 subjects showed broadband responses in their denoised data (Figure 9). All three subjects showed a higher rate of horizontal than vertical microsaccades in every stimulus condition (Figure 14), consistent with prior observations (33), but there was no systematic pattern in saccade frequency as a function of stimulus condition; for example, the stimulus condition with the most and with the fewest microsaccades differed across the 3 subjects. Moreover, the subject with the highest broadband SNR among these 3 (S6) had the lowest rate of microsaccades (~0.5 microsaccades / second). To test more directly whether microsaccades contributed to the measured broadband fields, we re-analyzed the data from these 3 subjects in two ways, either limited to only those epochs with microsaccades or only those epochs without microsaccades (Figure 14b). The broadband responses were evident in each subject in the epochs without microsaccades, indicating that this response is not entirely an artifact of microsaccades.

**Figure 14.**
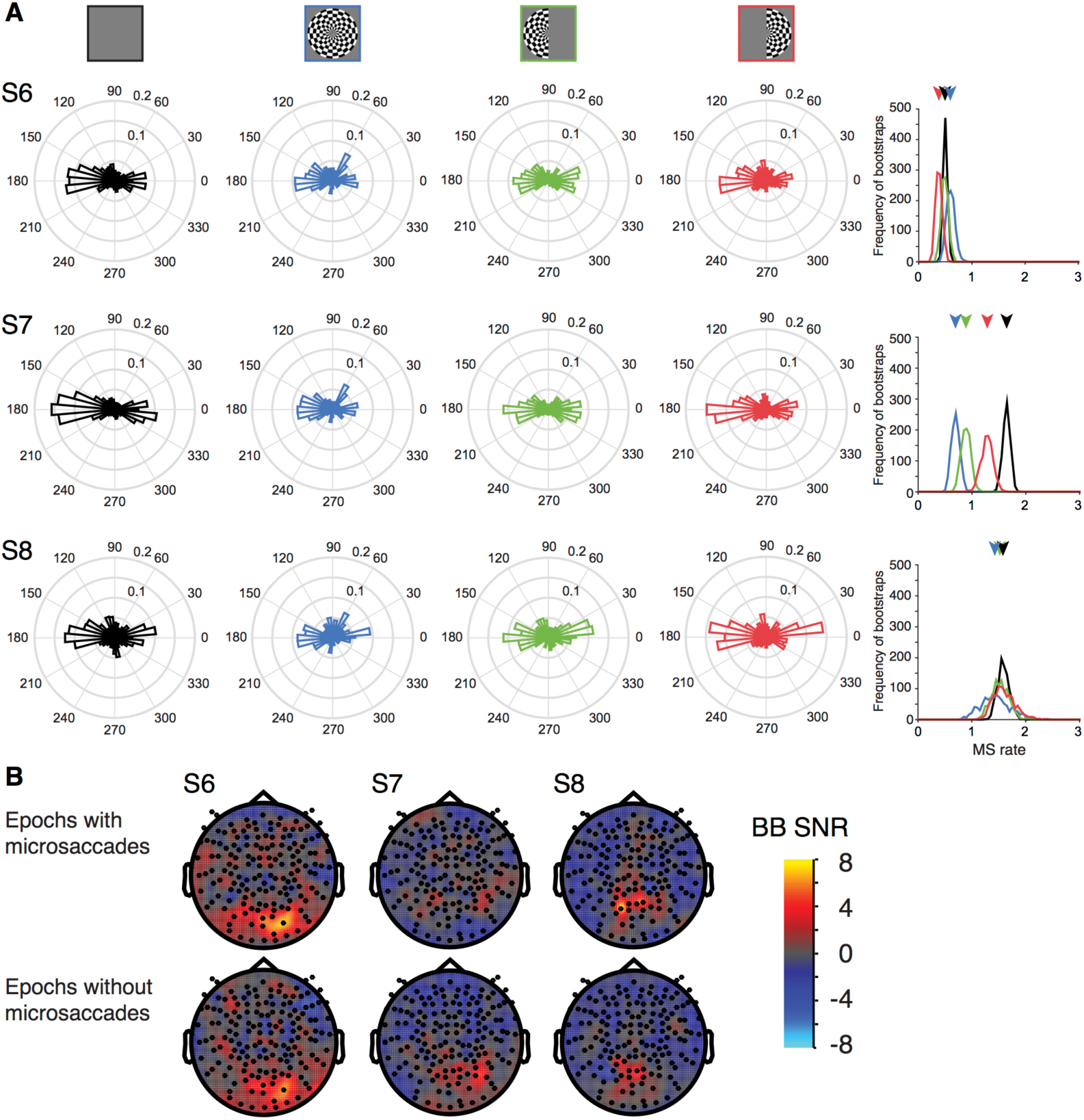
Microsaccades during experimental conditions. **(A)** The circular histograms show the frequency of microsaccades per 1-s epoch, binned by direction, for each of the 4 stimulus conditions (columns 1-4). The rows show data for 3 subjects. The last column shows the rate of microsaccades (per 1-s epoch) irrespective of direction, for each of the 4 stimulus conditions, bootstrapped 100 times over epochs. Arrows indicate the median rate for each condition. **(B)** Both-hemifield minus blank broadband SNR meshes limited to only those epochs with microsaccades (top row) or without microsaccades (bottom row). Made with *dfdMakeFigure14.m.*

## 3 Discussion

We separated the MEG signal into two components, one time-locked and one asynchronous with the stimulus. The stimulus-locked component was clearly visible in all subjects with minimal preprocessing. The asynchronous signal, spanning 60-150 Hz, was visible with little preprocessing in some subjects and the mean across subjects. However, the SNR was low compared to the stimulus-locked component. With our denoising algorithm, SNR more than doubled, resulting in reliable, spatially specific broadband signals in all individuals. We showed in a subset of subjects that the broadband signals could not be explained by systematic biases in the pattern of fixational eye movements, supporting the interpretation that the broadband fields arise from neural activity rather than artifacts associated with eye movements.

These results are *qualitatively* consistent with intracranial measurements (24). However, it has proven difficult to measure extracranial broadband signals arising from neural activity. Below, we discuss the significance of broadband responses, challenges in measuring them extracranially, and the generalizability of our denoising algorithm.

### 3.1 Significance of broadband responses

A century ago Berger and others described oscillations in surface EEG between 10 and 25 Hz (4, 34). More recently, using intracranial recordings, Crone and colleagues (35) reported power increases in higher frequencies (75-100 Hz) associated with motor movements. Subsequently, this high frequency response was interpreted as a broadband (not oscillatory) signal, thought to reflect increased neuronal activity, rather than increased synchrony (5, 6). In support of this interpretation, studies showed that broadband signals correlate with single (10) and multiunit spike rates (11), and are observed throughout cortex (Figure 15). Under some conditions, broadband is also correlated with the BOLD signal (13, 24). BOLD signals, however, are influenced by processes other than spiking (36-38); hence quantifying broadband responses from the same experimental paradigm studied with fMRI can help disentangle the relative contribution to the observed BOLD response from spiking versus other, non-spiking neural activity. Being able to reliably measure the broadband signal extracranially offers the opportunity to noninvasively measure neuronal spiking activity at a sub-second scale, complementing fMRI and oscillatory and time-locked (evoked) signals commonly measured with MEG and EEG.

**Figure 15.**
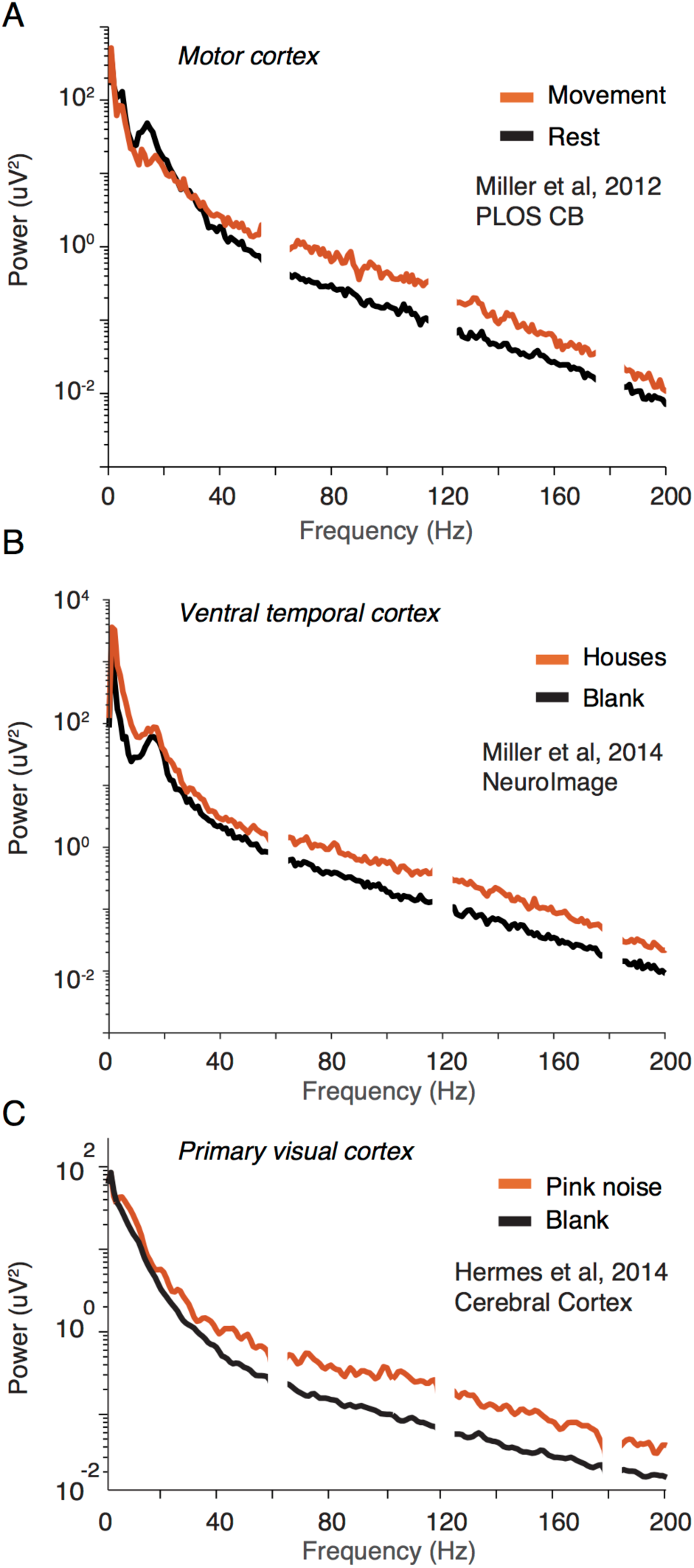
Broadband signals around the brain. ECoG studies have measured broadband power elevations associated with perception, movement, language, and cognition (7, 39-41) Examples of broadband field potentials from single ECoG electrodes in motor cortex **(A)**, ventral temporal cortex **(B)**, and primary visual cortex **(C)**. The power increases relative to baseline span at least 50 to 200 Hz. Adapted from (A) (42); (B) (7); (C) (43).

### 3.2 Prior measures of extracranial broadband and gamma band responses

#### Broadband vs. narrowband gamma

Several groups have distinguished broadband power increases from narrowband gamma oscillations (8, 11). Gamma oscillations are observed in visual cortex for some stimuli (e.g., high contrast gratings) (44, 45), with a peak frequency between 30 and 100 Hz and bandwidth of 10-20 Hz. The broadband response occurs in many brain areas for many types of stimuli, spanning at least 50-150 Hz, and can extend to lower and higher frequencies (6, 24). Robust gamma oscillations have been measured extracranially for grating stimuli (26, 27). These oscillations differ from the broadband fields reported here, which span a wider frequency range, have lower amplitude, and likely reflect asynchronous neural activity.

#### Multiple gamma peaks

Some extracranial studies have reported multiple distinct signals within the gamma band. For example, Wyart and Tallon-Baudry (46) and Vidal et. al. (47) measured MEG responses to gratings and bars, respectively. They reported power increases in two bands, from 45-65 Hz and 75-120 Hz. Both components were interpreted as oscillations arising from synchronous neural activity, and are likely different from the broadband signals reported here.

*Group averaged broadband.* Two MEG studies reported increases in high gamma power (60-140 Hz) during recall of visual stimuli (48, 49). These studies showed averages across subjects (22 or 24), so that it is not known whether there were reliable responses in individuals.

#### Motor cortex

High frequency signals (~65-100 Hz) have been shown from motor cortex measured extracranially (50, 51). This signal was most evident in group-averaged data and some but not all individuals, and within a relatively narrow band (~20-30 Hz wide). Ball *et al.* (50) noted that better methods for measuring high frequency broadband extracranially would help resolve whether individual differences were due to measurement limitations or the lack of high frequency brain signals in some subjects. Cheyne *et al.* (52) measured high gamma (65-80 Hz) with MEG over motor cortex in individual subjects, and speculated that these signals reflect cortico-basal ganglia loops, as the basal ganglia is known to produce narrowband oscillations peaked at 70-80 Hz.

### 3.3 Challenges in measuring extracranial broadband responses

#### Extracranial broadband signal strength is low

Although having high SNR after denoising, the MEG broadband signal was nonetheless small relative to baseline - about a 13% increase. Using nearly the identical stimulus, the broadband signal measured by ECoG was ~15 times larger (a ~190% increase) (24). The discrepancy was much smaller for the stimulus-locked signal (almost 8-fold increase with MEG vs. 21-fold with ECoG). Why are MEG broadband signals small? First, MEG sensors pool over a large area, so baseline power reflects activity from a large fraction of the brain, whereas visually driven broadband responses likely come from confined regions (53). In contrast, both baseline and visual responses in ECoG electrodes arise from the same cortical patch. Second, the amplitude depends not only on pooling area, but also phase coherence. If broadband signals arise from incoherent neural activity, and stimulus-locked signals from coherent (synchronous) activity, then the former will grow with the square-root of the number of sources, and the latter with the number of sources. Since MEG pools over much larger populations than ECoG, the ratio of incoherent signal strength (broadband) to coherent (stimulus-locked) will be much lower. This logic is supported by modeling (54) and empirical studies with intra- and extracranial measures, which found that the most coherent intracranial signals were best transmitted outside the head (55, 56).

#### Extracranial measurements contain multiple noise sources

Because extracranial broadband power is low, noise is a major impediment. In addition to neural noise (57), fixational eye movements (21), head muscle contraction (58), and environmental perturbations (22) produce noise measured by MEG and EEG sensors. Many of these noise sources are spectrally broad and hence particularly problematic when investigating neural broadband signals.

Although spike fields generated from eye movements can be mistaken for broadband neural activity (19), it is unlikely that our spatially-specific broadband measures were substantially contaminated by eye movement artifacts. This was confirmed by analyses of eye movement data, and the fact that middle posterior sensors where we observed broadband are not usually associated with MEG spike field artifacts (18). A second eye movement confound, the electromagnetic fields arising from movement of the retina-to-cornea dipole, causes low frequency artifacts (4-20 Hz; (59)) and therefore is unlikely to have affected our broadband measures (60-150 Hz).

Head muscles can also cause spectrally broadband contaminants (58), as can external noise sources, e.g., nearby electrical equipment. However, these noise sources are unlikely to be confined to occipital sensors and to co-vary with stimulus condition, and hence do not explain our broadband observations. Moreover, it is likely that these noise sources, if present, were included in our noise pool, and hence MEG Denoise would have reduced their effects.

### 3.4 MEG Denoise and other denoising algorithms

MEG Denoise uses PCA on a subset of sensors to remove noise. In principle, it can capture any noise source contributing to the noise pool, including environmental, oculomotor, muscular, and neural. This differs from algorithms designed to remove environmental noise. Hence MEG Denoise is complementary to these methods. We found that the most effective analysis was either MEG Denoise alone, or MEG Denoise following an environmental denoising algorithm. Our algorithm has much in common with ICA denoising (60), with some important differences. First, PCA, unlike ICA, ranks components by variance explained. Second, MEG Denoise explicitly identifies noise sensors. These features enable the algorithm to be fully automated, making it easy to denoise at the time scale of individual events (e.g., >1,000 one-second epochs). If the spatial pattern of the PCs vary over time, then deriving the components independently within short epochs is more effective (Figure 10).

To use MEG Denoise for other experimental designs, analyses, or scanners, one would need to change some input parameters. In addition to the experimental design matrix and data, required inputs include the experiment-specific functions to summarize the MEG responses. In our experiments, one function computed the stimulus-locked signal and was used to define the noise pool. For most of our analyses, a second function computed the broadband power to evaluate the results. In principle, one could use a single function to define the noise pool and evaluate the data (as we did for denoising the stimulus-locked signal). For other experiments, one might use a function that computes the amplitude or latency of an evoked response, or the power in a limited temporal frequency band, or any measure relevant to the experiment. Alternatively, one could run a separate localizer experiment to identify a pool of potential sensors of interest and a pool of noise sensors, and then manually enter the list of noise sensors to denoise the main experiment. There are several other optional inputs, such as the method to identify the noise pool, the accuracy metric (SNR/R^2^). Here, we used the defaults for all optional inputs.

### 3.5 Conclusion

Stimulus-driven broadband brain responses can be quantitatively characterized in individual subjects using a non-invasive method. Because we obtain high SNR measures from short experiments, the broadband signal can be used to address a wide range of scientific questions. Access to this signal opens a window for neuroscientists to study signals associated with neuronal spike rates non-invasively at a high temporal resolution in the living human brain.

## 4 Methods

### 4.1 Reproducible computation and code sharing

We adopt the view that the product of scientific research is not only the paper, but also data and software needed to reproduce the results (61-63). In the interest of reproducible research, both the analysis code and the MEG data for all results reported in this paper are publicly available via the Open Science Framework at the url https://osf.io/c59sh/ (doi 10.17605/OSF.IO/C59SH). All analyses were conducted in MATLAB. Figures 2-15 (except 3) can be reproduced by running scripts from the GitHub repository of the form *dfdMakeFigure4.m* (to reproduce Figure 4 from raw data), or the master script *dfdMakeAllFigures.m.*

### 4.2 Data acquisition

#### 4.2.1 Ethics Statement

The experimental protocol was in compliance with the safety guidelines for MEG research and was approved by the University Committee on Activities involving Human Subjects at New York University and by the ethics committee of the National Institute of Information and Communications Technology (NICT). Subjects provided written informed consent.

#### 4.2.2 Subjects

Eight subjects (five females), ages 20-42 years (M = 28.4 / SD = 6.7 years) with normal or corrected-to-normal vision participated in the NYU study. An additional 4 subjects (M = 27.0 / SD = 7.4 years) participated in the same experiment at Center for Information and Neural Networks (CiNet), National Institute of Information and Communications Technology (NICT) in Osaka, Japan. Display Stimuli were generated using MATLAB (MathWorks, MA) and PsychToolbox (64, 65) on a Macintosh computer. <u>NYU:</u> Images were presented using an InFocus LP850 projector (Texas Instruments, Warren, NJ) with a resolution of 1024 × 768 pixels and refresh rate of 60 Hz. Images were projected via a mirror onto a front-projection translucent screen at a distance of approximately 42 cm from the subject’s eyes (field of view: 22 deg × 22 deg). The display was calibrated with the use of a LS-100 luminance meter (Konica Minolta, Singapore) and gamma-corrected using a linearized lookup table. <u>CiNet</u>: The display parameters were similar, except that the projector was PT-DZ680 (Panasonic, Japan), with 800 × 600 resolution and 60 Hz, and 61 cm viewing distance.

#### 4.2.3 Stimuli

The stimuli were contrast-reversing dartboard patterns (12 square wave contrast reversals per second), windowed within either a half circle (left or right visual field) or full circle (bilateral visual field) aperture, with a diameter of 22 degrees at NYU (26 degrees at CiNet). Mean luminance gray (206 cd/m^2^ (NYU), 83 cd/m^2^ (CiNet)) was used as background color for the dartboards and was shown in the full field during blank trials between stimulus periods (Figure 1).

#### 4.2.4 Experimental design

One *run* consisted of six seconds flickering ‘on’ periods, alternated with six seconds ‘off’mean luminance periods, repeated 6 times (72 seconds). The order of the left-, right- or both-visual field apertures was random. There was a fixation dot in the middle of the screen throughout the run, switching between red and green at random intervals (averaging 3 seconds). The subjects were instructed to maintain fixation throughout the run and press a button every time the fixation dot changed color. The subjects were asked to minimize their blinking and head movements. After every 72-second run, there was a short break (typically 30-s to 1 minute). Each subject participated in 15 runs.

#### 4.2.5 MEG signal acquisition

Data for the main experiment were acquired continuously with a whole head Yokogawa MEG system (Kanazawa Institute of Technology, Japan) containing 157 axial gradiometer sensors to measure brain activity and 3 orthogonally-oriented reference magnetometers located in the dewar but away from the brain area, used to measure environmental noise. The magnetic fields were sampled at 1000 Hz and were filtered during acquisition between 1 Hz (high pass) and 200 Hz (low pass).

In a subset of subjects (S6-S8), eye movements were recorded by an EyeLink 1000 (SR Research Ltd., Osgoode, ON, Canada). Right eye position data were continuously recorded at a rate of 1000 Hz. Calibration and validation of the eye position was conducted by having the subject saccade to locations on a 5-point grid. Triggers sent from the presentation computer were recorded by the EyeLink acquisition computer. The same triggers were recorded simultaneously by the MEG data acquisition computer, allowing for synchronization between the eye-tracking recording and MEG recording.

The 4 data sets acquired with an Elekta Neuromag at CiNet and were pre-processed in MATLAB (MathWorks, MA, USA) using the identical code and procedure. The CiNet data were acquired as 102 pairs of planar gradiometer signals (204 sensors). Data were analyzed from each of the 204 gradiometers separately and paired into 102 locations for mesh visualization (e.g., the broadband signal-to-noise-ratio for sensor 121 and 122 out of 204 would be averaged to show one signal-to-noise-ratio in the position of sensor 61 out of 102).

### 4.3 Data analysis

#### 4.3.1 MEG preprocessing

For some analyses, data were environmentally denoised using published algorithms prior to any further analysis. This enabled us to compare data denoised with our new algorithm alone, or with our new algorithm following environmental denoising. For the NYU data, we used either of two algorithms. One was the continuously adjusted least-square method (CALM; (29), applied to data with a block length of 20 seconds (20,000 time samples). The second algorithm was time-shifted principal component analysis (TSPCA; (30), with a block length of 20 seconds and shifts of up to +/- 100 ms in 1 ms steps. For the CiNet data, the environmental denoising algorithm was temporal signal space separation (‘tSSS’) (with default parameters, e.g. inside and outside expansion orders of 8 and 3, respectively; 80 inside and 15 outside harmonic terms; correlation limit of 0.98).

The FieldTrip toolbox (66) was used to read the data files (either environmentally-denoised or raw). For all subsequent analyses, custom code was written in MATLAB. Using either the environmentally-denoised data or raw data, the signals were divided into short epochs. Each stimulus type (left-, right-, or both-hemifield, or blank) was presented in 6-s blocks, and these blocks were divided into 6 non-overlapping 1-s epochs. We discarded the first epoch of each 6-s block to avoid the transient response associated with the change in stimulus. After epoching the data, we used a simple algorithm to detect outliers. We first defined a ‘data block’as the 1-s time series from one epoch for one sensor. So a typical experiment consisted of ~170,000 data blocks (157 sensors × 1080 1-s epochs). We computed the standard deviation of the time series within each data block, and labeled a block as ‘bad’ if its standard deviation was more than 20 times smaller or 20 times larger than the median standard deviation across all data blocks. The time series for bad data blocks were replaced by the time series spatially interpolated across nearby sensor (weighting sensors inversely with the distance). Further, if more than 20% of data blocks were labeled bad for any sensor, then we removed the entire sensor from analysis, and if more than 20% of data blocks were bad for any epoch, then we removed the entire epoch from analysis. Typically, two to seven sensors and 2%-4% of the epochs were removed per session for the NYU data. For the CiNet datasets, almost no sensors or epochs were removed (one sensor and one epoch across all data sets). These preprocessing steps were implemented with *dfdPreprocessData.*m.

#### 4.3.2 Computation of stimulus-locked and broadband responses

Data were summarized as two values per sensor and per epoch: a stimulus-locked and a broadband power value. These calculations were done by first computing the Fourier transform of the time series within each epoch (Figure 16A,B).

The stimulus-locked signal was then defined as the amplitude at the stimulus-locked frequency (12 Hz). The broadband response was computed as the geometric mean of the power across frequencies within the range of 60-150 Hz, excluding multiples of the stimulus-locked frequency (see also Figure 16AB). The geometric mean is the exponential of the average of the log of the signal. We averaged in the log domain because log power is better approximated by a normal distribution than is power, which is highly skewed. These two calculations converted the MEG measurements into a broadband and a stimulus-locked summary metric, each sampled once per second (Figure 16C). The two summary metrics were computed by the functions *getstimlocked.m* and *getbroadband.m.*

**Figure 16.**
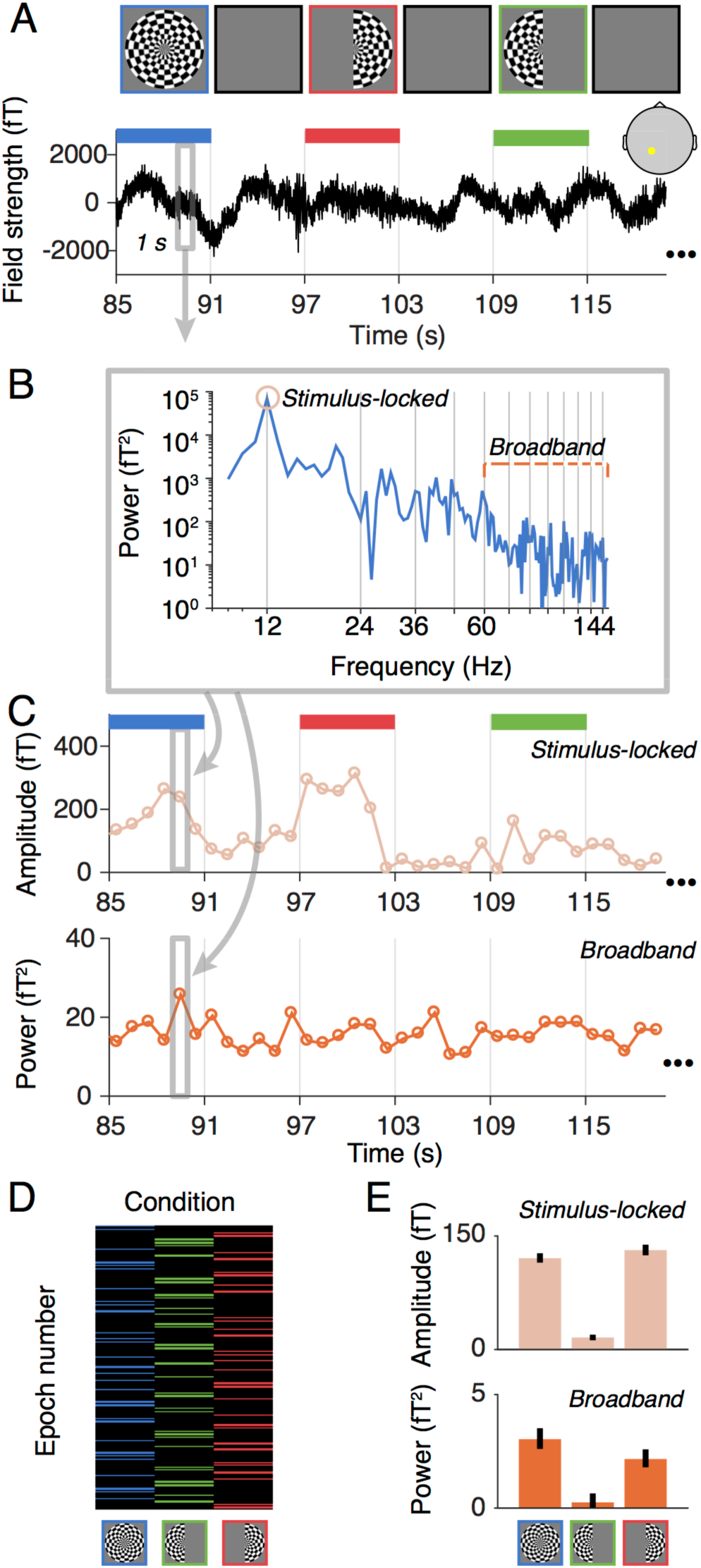
Data analysis without denoising. **(A)** The time series for each sensor were epoched into non-overlapping one-second periods. **(B,C)** The time series in each epoch was fast Fourier transformed and then summarized as two values, a stimulus-locked value (amplitude of the fast Fourier component at the stimulus frequency), and a broadband value (mean of the log power of all frequencies from 60-150 Hz, excluding those within +/− 1 Hz of stimulus harmonics). **(D)** The summary of conditions is shown as a matrix, where each column corresponds to one of the three stimulus conditions, and the number of rows is equal to the total number of epochs across the session. Rows with no color are blank epochs. (**E**) Summary metrics were computed separately for the stimulus-locked values and broadband measures, yielding three measures per sensor per data type. The summary metric was the mean across condition minus the mean across blanks, bootstrapped 1000 times. The bar plots show the median and the 68% confidence interval based on 1000 bootstraps.

We then bootstrapped across epochs to compute confidence intervals on the signal estimates (per sensor and per condition). For each of 1000 bootstraps, we sampled *n* epochs with replacement, where *n* is the total number of epochs in the experiment. We then computed the average response across epochs for each stimulus condition, minus the average across blank epochs. This provided one summary measure for each of the three stimulus conditions and each of the two dependent measures (broadband and stimulus-locked) for each of the 1000 bootstraps. Finally, we took the median across bootstraps as the estimate of signal and half of the 68% confidence interval across bootstraps as the estimate of the noise (Figure 16D,E). For some analyses, the ratio of these values was defined as the signal-to-noise ratio (SNR).

#### 4.3.3 MEG Denoise Algorithm

Extracranial measurements like MEG have multiple global noise sources and a relatively low signal-to-noise ratio compared to intracranial measures, especially for high frequency signals. In order to increase the signal-to-noise ratio, we developed a denoising technique that helps reveal the broadband signal of interest. A denoising algorithm developed for fMRI (‘GLMdenoise’; (1)) was adapted for MEG to project out noise from the data for each epoch in each sensor. The logic behind the algorithm is that many sources of noise are global, and therefore spread across sensors. The algorithm identifies sensors that have no stimulus-related response (the ‘noise pool’), and uses these sensors to define noise components. The noise components are then projected out from all sensor time series in each epoch.

##### Noise pool selection

The noise pool was defined as the 75 (NYU) or 100 (CiNet) sensors with the lowest stimulus-locked SNR across conditions. The SNR was computed by (a) dividing the median response across bootstraps by the variability across bootstraps (half of the 68% confidence interval) for each condition, and (b) taking the maximum of the three values (corresponding to the three stimulus conditions) for each sensor.

We used the stimulus-locked signal to identify the noise pool because this signal had a very high SNR, and could easily by measured prior to running our denoise algorithm, and because we assumed (and confirmed by inspection) that sensors with broadband responses also had stimulus-locked responses.

For most subjects, most of the sensors in the noise pool were located over the front of the head (see for example Figure 4A).

##### Filtering of time series

As described above, the broadband summary metric was derived from power at a limited range of temporal frequencies (60-150 Hz, excluding multiples of the stimulus frequency). After defining the noise pool, the time series of all sensors in all epochs were filtered to remove signal at all frequencies not used to compute the broadband signal. Hence the remaining time series contained power only at frequencies defining the signal of interest. This step was important because the noise pool, though selected for a low stimulus-locked SNR, could nonetheless have contained a small, residual stimulus-locked signal. This residual signal would have been correlated with the experimental design (larger when stimuli were present than absent) and hence projecting it out of the data could have caused a systematic bias (see the script *denoisingProjectingInVariance.m).*

##### PCA

Following filtering, the next step in the algorithm was principal component analysis (PCA). This identified the common components of the time series across the sensors in the noise pool. PCA was computed separately for each 1-s epoch. This means that denoising occurred at the same temporal scale (1 second) as the computation of the summary metrics. This differs from some denoising algorithms, in which noise regressors are identified over a much longer time period, e.g., several minutes (60). Denoising at a short-time scale can be advantageous if the spatial pattern of the noise responses is not consistent across the entire experiment. As a control comparison, we also ran our algorithm by identifying PC time series on the entire duration of the experiment (~20 minutes) rather than epoch by epoch. (See Results, ‘Control analyses for MEG Denoise algorithm’.)

##### Projecting out PCA components

The first one to ten principal components (PCs) in each epoch were projected out of the time series for all sensors, using linear regression. This resulted in ten new data sets: One with PC 1 projected out, one with PC 1 and 2 projected out, etc. up to 10 PCs projected out. After projecting out the noise components, we summarized the data into a stimulus-locked and broadband component as described in Figure 16.

#### 4.3.4 Statistical comparisons

To assess the effect of the MEG Denoise algorithm on the broadband SNR, we compared the broadband SNR after applying MEG Denoise to the broadband SNR either without denoising or after applying other denoising algorithms. To make these comparisons, we first identified 10 sensors of interest from each subject. These sensors of interest were the 10 with the highest SNR in any of the three stimulus conditions, either before or after denoising, excluding sensors from the noise pool. For each of the three stimulus conditions, we then took the average SNR from these 10 sensors without denoising or after applying MEG Denoise or another denoising algorithm. Finally, we conducted two-tailed t-tests, paired by subject (n=8), between the broadband SNR after MEG Denoise to the broadband SNR without denoising (or with another algorithm). The t-tests were conducted separately for each of the three stimulus conditions (both-hemifield, left-hemifield, and right-hemifield).

#### 4.3.5 Control analyses

To investigate the validity of our algorithm, we ran multiple control analyses. In particular, it is important to rule out the possibility that the denoising algorithm produces significant results even when the data contains no sensible signal. To test this, we compared the difference in SNR of denoised data with the following controls: (1) phase-scrambling the PC time series, and (2) using all sensors to define the noise with PCA rather than only a subset of sensors that have little to no stimulus-locked signal. We also assessed the effect of identifying and projecting out PC time series equal in length to the entire experiment (~20 minutes), rather than PC time series matched in length to our analysis epochs (1-s). This comparison tested the assumption that denoising in shorter epochs was advantageous, possibly due to the pattern of noise sources differing over the course of the experiment.

#### 4.3.6 Eye tracking analysis

Since an increase in microsaccade rate can induce broadband spectral components in extracranial measurements such as EEG or MEG (<u>21,</u> 59), we checked in three NYU subjects (S6-S8) whether there was a difference in rate between the ‘off’ baseline periods and ‘on’ stimulus periods, and within the three stimulus (both-, right-, left-hemifield) conditions. Microsaccades were identified as changes in position with above a relative velocity threshold (6°/s) and a minimum duration of 6 ms, as reported in Engbert & Mergenthaler (67) to analyze rate and direction of microsaccades as well as separating MEG data into epochs that did and did not contain microsaccades.

## Acknowledgments

This study was supported NEI grant R00-EY022116 (J.W.), and NIMH grant R01-MH111417 (J.W.).

## Supplementary Figures

**Supplementary Figure 1.**
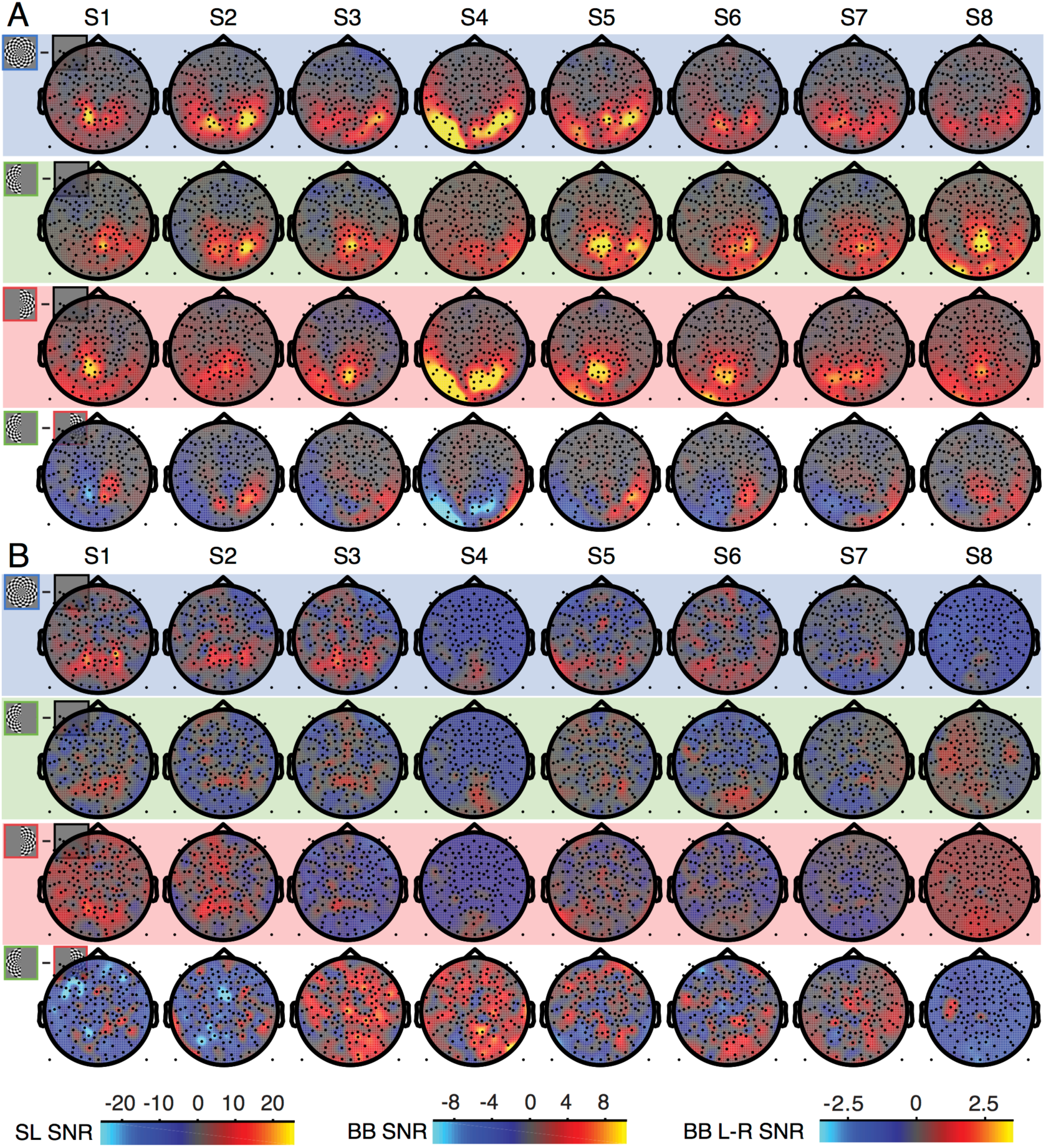
Topographic maps of stimulus-locked and broadband SNR in individual subjects before denoising. **(A)** Stimulus-locked SNR for the both-, left-, right-, and left minus right-hemifield stimulus, without denoising. Rows 1-4 use the SL SNR color bar. **(B)** Broadband SNR before denoising. Rows 1-3 use the BB SNR color bar, and the 4^th^ row uses the BB L-R SNR color bar.

**Supplementary Figure 2.**
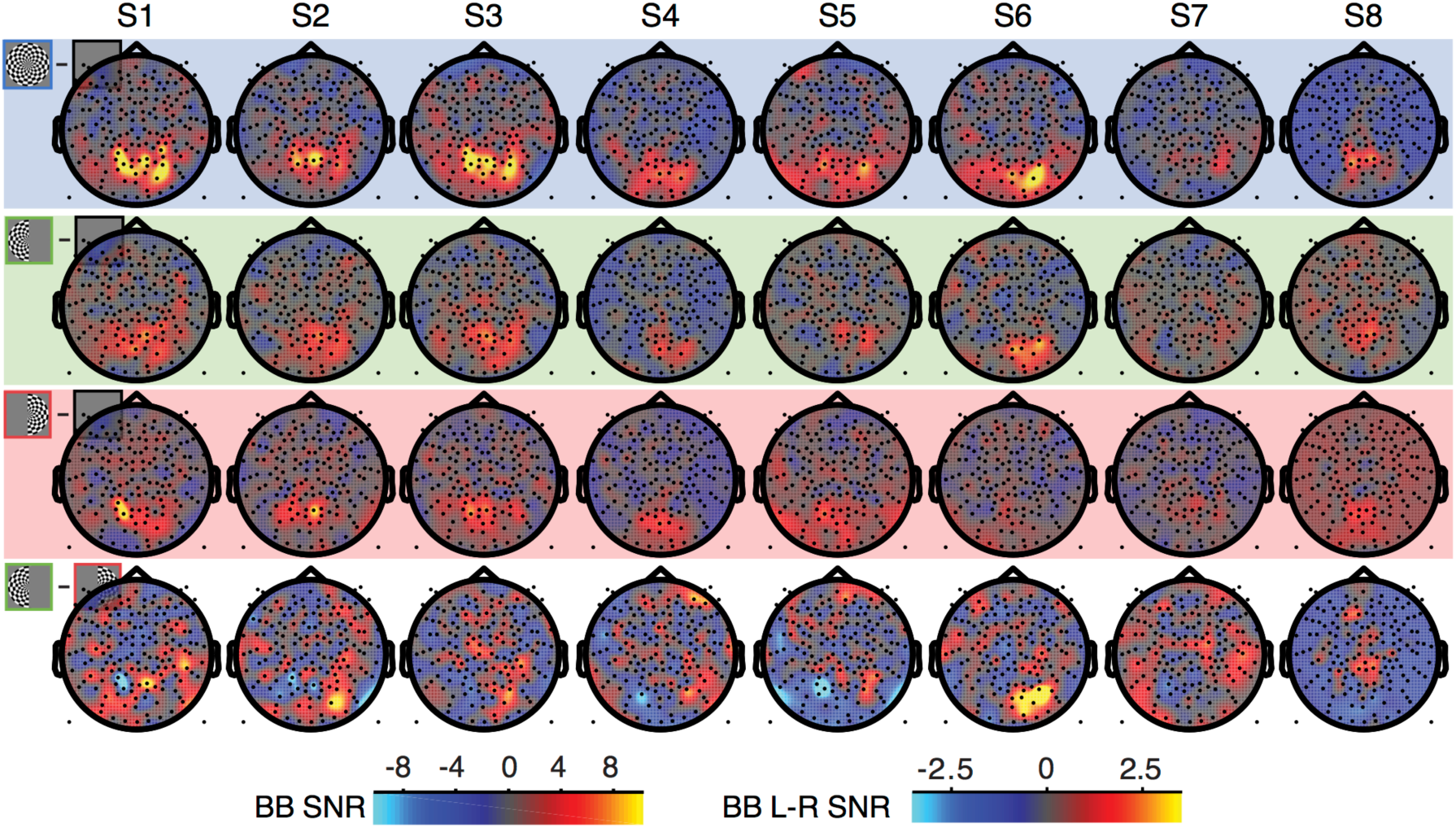
Topographic maps of broadband SNR in individual subjects after denoising. **(A)** Head plots show the stimulus-locked SNR for the both-, left-, right- and left minus right-hemifield stimulus, after denoising. Rows 1-3 use the BB SNR color bar, and the 4^th^ row uses the BB L-R SNR color bar.

